# Diverse Conformational Ensembles Define the Shared Folding-Allosteric Landscapes of Protein Kinases

**DOI:** 10.1101/2025.08.13.670202

**Authors:** Dhruv Kumar Chaurasiya, Athi N. Naganathan

## Abstract

Sequence variation across members of an enzyme family contributes to diverse ensemble behaviors, which subtly influence substrate affinity, selectivity and regulation. A classic example is the family of eukaryotic protein kinases (EPKs), which regulate numerous cellular processes and serve as important drug targets. Here, we dissect the consequences of sequence variation on the folding-conformational landscapes by performing a meta-analysis of 274 EPKs through a structure-based statistical mechanical framework. We find that EPKs populate several partially structured states in their native ensemble with a hierarchy of structural order in the N-terminal lobe that is critical for catalysis and activation. Despite this, the unfolding mechanism is uniquely conserved across the majority of kinases, with the N-terminal lobe unfolding first. Kinase activation modulates the local stability and thermodynamic connectivity in a non-conserved manner and across the entire structure, due to the strong coupling between the active site residues to distant sites, including the established allosteric pockets. We further show how activation drives the Abl kinase ensemble towards a more folded and thermodynamically coupled system in a graded manner. Our work uncovers the thermodynamic design principles of kinases with insights into allostery, while shedding light on the extents to which ensemble behaviors are impacted by sequence variations in paralogs.

## Introduction

Kinases constitute a family of structurally similar enzymes responsible for catalyzing the transfer of phosphate groups from adenosine triphosphate (ATP) to serine, threonine, or tyrosine residues on target proteins. This phosphorylation event is tightly regulated and is a cornerstone of intracellular signaling, influencing numerous cellular processes such as cell cycle progression, metabolism, differentiation, and apoptosis.^1–4^ Kinases can be classified into distinct subfamilies based on substrate specificity, including serine/threonine kinases, tyrosine kinases, and dual-specificity kinases,^5^ each playing a critical role in maintaining cellular homeostasis. Disruptions in kinase function - either through mutations or over-expression - are implicated in various pathologies, particularly cancer.^6^ Accordingly, kinases are much sought after therapeutic targets and identifying inhibitors that are selective to specific kinases is an active area of research.^6–10^

Structurally, kinases have two lobes, termed the N-terminal (N_t_; with five β-sheets an α-helix) and the C-terminal (C_t_; with six α-helices) lobes (Figure 1a).^11^ The active site where ATP, Mg^2+^ and the substrate bind is located at the interface or a cleft between the two lobes. There are multiple conserved structural motifs whose conformational features determine the state of kinases, i.e. active or inactive, and their functionality (Figure 1b). The Gly-rich loop (GxGxxG or the G/P-loop or the phosphate-binding loop) is located between the strands β_1_ and β_2_ and is responsible for positioning the γ-phosphate of ATP for catalysis. The αC-helix is a dynamic motif whose relative conformation along with the activation loop (ALN) is critical for kinase activation mechanisms.^12–16^ The catalytic loop between αE and β_6_ contains the H/Y-RD and contributes a large fraction of the catalytic machinery, while the activation loop between β_7_ and αF loop harbors the DFG motif with the Aspartate (D) critical for recognizing the ATP-bound Mg^2+^ ion. The catalytic loop backbone is rigid even in the apo-form as well as in the inactive state. However, the activation loop together with the DFG-motif and the αC-helix undergo dramatic and coupled structural re-arrangements in transitioning from inactive to the active state. In the inactive kinase, the DFG-motif exhibits a ‘DFG-out’ conformation with the activation loop folded back into the structure and occluding substrate binding and far from the αC-helix. In the active kinase, the αC-helix undergoes a rigid-body rotation towards the substrate binding cleft, with the activation loop adopting an extended surface for substrate binding.^17,18^ In addition, the DFG-motif now adopts a ‘DFG-in’ conformation with the aspartate positioning itself in the ATP-binding pocket. These structural arrangements further bring together multiple residues in proximity completing the ‘regulatory-’ (R-) and ‘catalytic’-spines (C-spine).^22,23^

**Figure 1.**
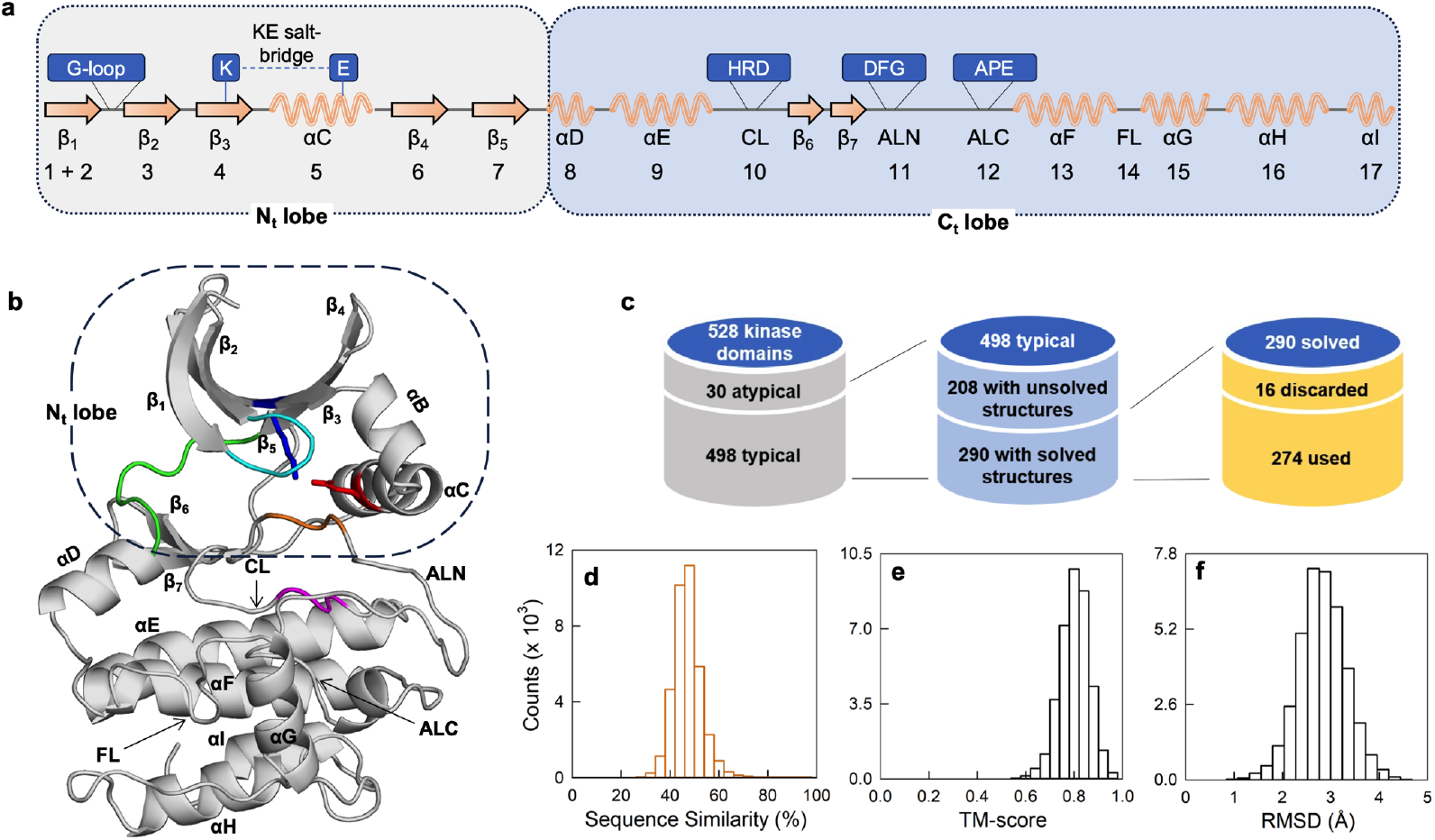
Structural architecture of kinases and database construction. (a) Schematic one-dimensional representation of kinase structure with conserved sub-domains. α-helices and β-strands are represented as springs and arrows, respectively. Regions marked as CL, ALN, ALC, and FL are catalytic, N-terminal activation loop, C-terminal activation loop, and F loop, respectively. Conserved regions such as glycine-rich loop (G-loop), lysine (K)-glutamate (E) salt-bridge, histidine (tyrosine in AGC family)-arginine-aspartate (H/YRD), aspartate-phenylalanine-glycine (DFG) and alanine-proline-glutamate (APE) motifs are also highlighted. The numbers below secondary structure cartoons represent the sub-domains according to Dunbrack’s classification^19^. Note that αB helix is present only in kinases belonging to AGC, CAMK and ‘Other’ family. (b) The structure of catalytic domain of protein kinase A (PDB id: 4WB8) with marked conserved structural domains. The Gly-rich loop (cyan), catalytic Y/HRD motif (magenta), activating DFG motif (orange), hinge loop (green) along with the conserved Lys of β_3_ (blue sticks) and Glu of αC (red sticks) are shown separately. (c) Selection criteria for construction of the kinase database. (d) Distribution of pairwise sequence similarity amongst the 274 kinases calculated using Clustal Ω^20^ with a mean similarity percentage of 47.3 ± 6.3 %. (e, f) Distribution of pairwise structural similarity calculated using mTM-align^21^ that yields a mean TM-score of 0.80 ± 0.06 and a mean RMSD of 2.8 ± 0.5 Å.

The known sequence diversity of kinases arises primarily from variation in sequence across the protein, while the catalytic and residues or motifs involved in activation are naturally conserved.^24–27^ This raises questions on the mechanisms through which selectivity in both phosphorylation and regulation is achieved. It is likely that a large fraction of protein residues acts towards defining a function which includes substrate selectivity, affinity and regulation, as reported in many systems.^28–31^ How different then are the intra-molecular interaction networks in kinases and hence their thermodynamic architecture? Does a conservation of the overall topology and fold naturally lead to similar native ensembles and (un)folding mechanisms? Does sequence diversity at regions distant from the active site promote varied populations of partially structured states despite the similar architecture? Do they respond differently to activation? In other words, does the diversity in the intra-molecular interaction network lead to different coupling extents across the structure? Is it possible to quantify them and derive insights into allostery?

The sheer number of eukaryotic kinases (>500) and their sizes (>250 residues) rule out MD simulations as a tool to explore these specific questions on a large scale. On the other hand, a structure-based statistical thermodynamic treatment of the kinase conformational landscapes could reveal insights into conserved and non-conserved attributes of kinases, folding mechanisms, and potential differences in allosteric connectivity. In this work, we employ the predictive and successful Wako-Saitô-Muñoz-Eaton (WSME) model of protein folding^32–34^ to extract conserved thermodynamic design principles underlying the native ensemble of 274 eukaryotic kinases (Figure 1c). Despite the sequence diversity (Figure 1d), kinases follow a conserved unfolding mechanism, but with diverse native ensembles, and large differences in local stabilities, which naturally lead to non-conserved sensitivity to perturbations. We find that most of the known allosteric pockets are thermodynamically coupled to the orthosteric site but to different extents across kinases, highlighting how sequence patterning around the active site can lead to dramatic functional outcomes despite similar three-dimensional structures (Figure 1e, 1f). We finally validate these predictions via a detailed analysis of the native conformational landscape of Abl1 kinase, which displays graded thermodynamic coupling patterns in different functional states.

## Methods

### Kinase database

Structures of human catalytic kinase domains were downloaded from Kincore.^35^ These were identified using the list of kinase genes available in UniProtKB’s pkinfam (https://ftp.uniprot.org/pub/databases/uniprot/current_release/knowledgebase/complete/docs/pkinfam.txt) that contain 484 typical and 30 atypical human kinases (Supporting Table S1). Fourteen of the genes out of 484 typical kinases translate to two catalytic domains, resulting in a total of 498 typical kinase domains (Figure 1c). Atypical kinases are not included in the database due to their large structural variability. From among the remaining 498 domains, only 290 have solved structures in the RCSB PDB database (Table S1). A representative structure with the highest resolution for each catalytic domain of kinase (referred to as kinase from now) was then downloaded with sequence boundaries defined by UniProtKB and renumbered from one. For cases where multiple structures of a kinase domain are available in the RCSB PDB database, the highest resolution structure with minimum mutations and missing residues was downloaded. Since most kinase structures in Kincore are in their active conformation (∼4300), structures in the active state are preferred over their inactive state, and only one chain of the asymmetric unit in the structure file is used. Thus, the final database has 202 kinase structures in the active state, while the remaining 88 are inactive.

To model missing atoms or residues, the kinases are remodeled using the comparative modeling module of RoseTTAFold,^36,37^ with its respective PDB structure as the template. Twelve kinase structures (Table S2) are excluded due to a large number of missing or deleted residues. Additionally, four highly dissimilar structures (Figure S1, Table S2) with a mean template modeling score (TM-score) less than 0.7 are also excluded from the database, reducing the total number of kinase structures to 274. Of these 274 kinases, only 104 have structures in both active and inactive conformations, resulting in 104 active-inactive pairs. The complementary high-resolution structures for each pair were downloaded and passed through RoseTTAFold to remodel missing atoms and residues.

### Wako–Saitô–Muñoz–Eaton (WSME) model

The WSME model is a structure-based statistical mechanical model which is extensively discussed in previous works.^34,38^ In this study, we employ the block-version of the WSME model (bWSME), wherein a group of residues, termed blocks, act as independent folding units. Each block can be either folded (represented by *1*) or unfolded (*0*), and the ensemble is represented by microstates with only a single stretch of folded blocks (single sequence approximation, SSA), two stretches of folded blocks separated by an island of unfolded blocks (double sequence approximation, DSA), and DSA with interactions between the folded stretches if they do interact in the native structure (DSA with loop, DSAw/L). This approximation generates a large ensemble of microstates which can be envisioned as strings of *0*s and *1*s. The predictions of the bWSME model are quite robust and have already been validated on multiple systems. In this study, we employ a block size of 3, three consecutive residues correspond to one block while ensuring no block occupies two different secondary structure regions, for all the kinases.

The total partition function (*Z*) of the bWSME model is calculated as

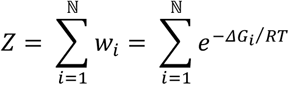

where ℕ is the total number of microstates (consisting of SSA, DSA, DSAw/L sub-ensembles), *R* is the gas constant, *T* is the temperature (310 K), and *w*_*i*_ is the statistical weight of state ‘*i*’. The free energy of each microstate (*m, n*) having a folded residues between and including residues *m* and *n* is

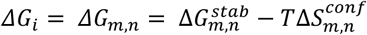

The stabilization free energy, 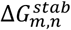, includes contributions from van der Waals interactions (defined by vdW interaction energy (ξ) for the contacts present within a 5 Å cut-off, including the nearest neighbors), contact-based implicit solvation free energy ΔG_solv_ (defined using the heat capacity change per native contact 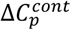, which is fixed to -0.36 J mol^-1^ K^-1^ per native contact^38^), and electrostatic interactions at pH 7.0 without any distance cut-off (through the Debye-Hückel formalism^38^). The entropic penalty for fixing the blocks in the folded conformation for any microstate defined by (*m, n*) is given by

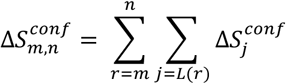

where, 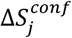 is the entropic penalty for fixing the residue *j* in the folded conformation (assigned a value of -14.5 J mol^-1^ K^-1^ per residue), while *L*(*r*) includes the set of residues within the block ‘*r*’. An excess entropic penalty (ΔΔ*S*) of -6.1 J mol^-1^ K^-1^ per residue is additionally assigned to glycine and STRIDE-identified coil residues,^39^ owing to their higher flexibility.^40^ The entropic penalty for proline residues is fixed to 0 J mol^-1^ K^-1^ due to their conformational rigidity. Partial partition functions are calculated by grouping microstates with a specific number of folded structured blocks, from which one-dimensional free energy profiles and two-dimensional free-energy surfaces are generated.

The probability (*p*_*x*_) and stability *ΔG*_*s,x*_ of a block *x* is calculated as,

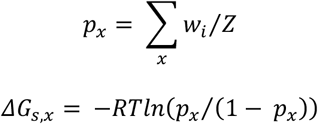

where the summation runs over all microstates in which block *x* is folded. All calculations are performed under iso-stability conditions of -30 kJ mol^-1^; this is achieved by tuning the ξ for every kinase while keeping all other parameters constant, unless mentioned otherwise. For the inactive-active kinase pairs, the ξ of the latter state was employed. All the calculations are performed without ligands, and information from only monomeric polypeptides is used to generate the ensemble of states with their relative statistical weights.

Structure-based sequence alignments are used to demark the boundary between the N and C terminal sub-domain, hereafter referred to as N_t_ and C_t_ lobe, wherein the start of the αD helix marks the start of the C_t_ lobe. The lobe stability (⟨ΔG_s_⟩) is then calculated as,

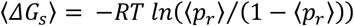

where ⟨*p*_r_⟩ is the mean folding probability of the residues that constitute the lobe. The folding probability of a residue *i* at a given point along the reaction coordinate, the number of structured blocks *n*, is calculated as,

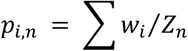

where the summation runs over all the microstates in which residue *i* is structured, and *Z*_*n*_ is the partial partition function associated with *n* structured blocks. Contact densities are calculated by taking the mean of interactions generated from the contact-map either globally or according to the definition of N_t_ and C_t_ lobes. A contact is considered non-local if the sequence separation between the interacting residues is greater than two.

### Thermodynamic coupling free energies

The bWSME model ensemble can also be partitioned into four sub-ensembles for every residue pair (*i, j*) arising from the conditional folded status of the residues involved in the collection of microstates: a sub-ensemble in which both *i* and *j* are folded 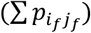, both *i* and *j* unfolded 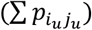, only *i* folded but not 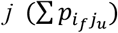, and only *j* folded but not 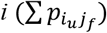.^41^ Here, the summation runs over the individual probabilities of microstates according to the proposed partitioning. These sub-ensembles are employed to calculate positive, negative, and effective thermodynamic coupling free energies from,

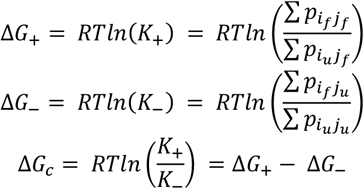

Here *K*_+_ and *K*_%_ represent the equilibrium constants for positive and negative coupling, respectively.

### Alanine-scan

Computational alanine-scanning mutations were carried out on a collection of kinases (PRKACA from AGC, PIM1 from CAMK, CSNK1A1 from CK1, MAPK1 from CMGC, NEK1 from NEK, MAP4K1 from STE, RIPK2 from TKL, ABL1 from TYR and ULK3 from ‘Other’ family) wherein all the residues were mutated to alanine employing PyMol,^42^ except for glycine, proline and alanine. The mutant (*mut*) structures were then fed into the model which re-calculates all interactions (including vdW and electrostatics) and predicts free-energy profiles and coupling free energies, employing identical parameters as the wild-type (*wt*). The positive coupling free energy (Δ*G*_+_) is then used for generating the differential coupling matrix (DCM) as ΔΔ*G*_+_ = Δ*G*_+,*mut*_ - Δ*G*_+,*wt*_. The distance-dependence effect of the mutation is calculated by averaging the ΔΔ*G*_+_ to obtain differential coupling indices (DCI) as a *N*×*1* vector, with N being the residue number; DCI then plotted as a function of C_*α*_ distance from the mutated residue (*d*), fit to a function of the form 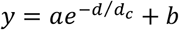 to extract the extent to which distant residues are affected, i.e. the coupling distance *d*_*c*_ in Å. In the above equation *a* is the amplitude in kJ mol^-1^ and *b* is the shift (in kJ mol^-1^).

## Results and Discussion

### Kinases sample multiple sub-states in their native ensemble

The bWSME model employs a binary description of folded status of individual residues or blocks (a set of consecutive residues) to generate a large ensemble of microstates of conformational states (Figure 2a and Methods).^38^ The statistical weight (*w*) of every microstate is dependent on the balance between stabilization (free-)energetic terms (Δ*G*^*stab*^ that includes contributions from van der Waals interaction energy, electrostatics, and temperature-dependent implicit solvation), and conformational entropy (Δ*S*^*conf*^), with the latter accounting for sequence-structure contributions. The stabilization free energies are derived from the native structure, and hence the model is Gō-like in its construction. The partition function (*Z*) is calculated from the statistical weights of the microstates, while partial partition functions can be employed to construct one-dimensional free energy profiles and even two-dimensional free energy surfaces as a function of the number of structured blocks. In addition to block-level folding probabilities or local stabilities, it is possible to calculate pair-wise correlation terms reporting on higher-order interactions termed the effective coupling free energies (Δ*G*_*c*_),^41^ derived from positive and negative coupling free energies (Δ*G*_+_ and Δ*G*_−_, respectively, in Figure 2a).

**Figure 2.**
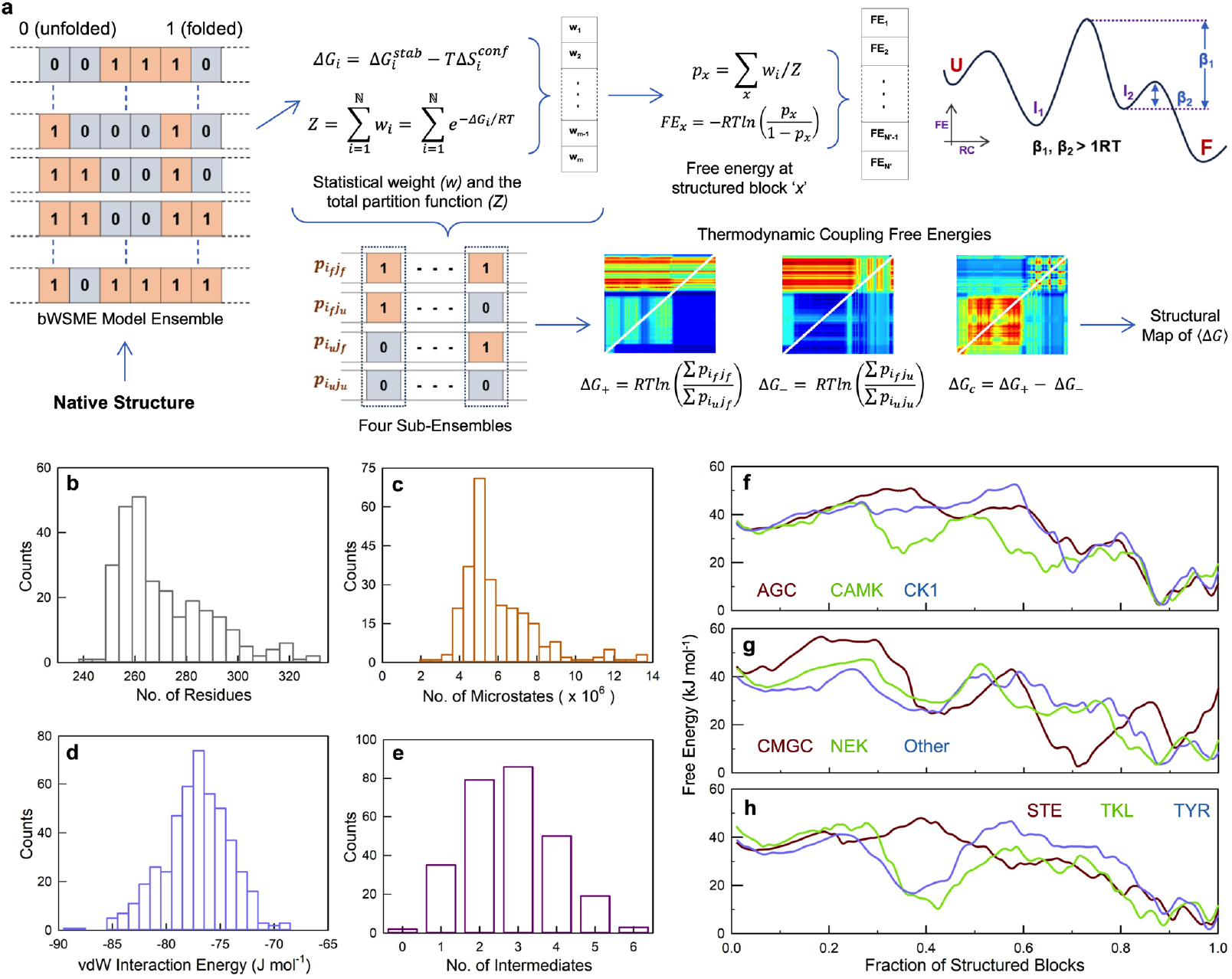
The bWSME model and the conformational landscape of kinases. (a) Schematic representation of the approach employed for calculating free energy (FE) profiles and thermodynamic coupling free energies from the bWSME model with just the native structure as an input (see Methods for details). In the representative free energy profile (top right), U, F, I_1,_ and I_2_ denote the unfolded, folded, intermediates 1 and 2, respectively. The barrier height (β) is defined as the energy difference between the maximum and minimum points on the FE profile. A minimum is considered an intermediate state only if both the unfolding and folding barrier height (β_1_ and β_2_ respectively) associated with that minimum exceeds 1*RT*. (b) The distribution of the number of residues in kinases. Majority of kinases are ∼260 residues in length, except for those belonging to CK1 and CMGC family which have ∼290 residues. (c) The distribution of the number of microstates in the bWSME model. (d) The distribution of vdW interaction energy (ξ) with a mean ξ of -76.5 ± 2.9 J mol^-1^ per native contact. (e) Distribution of the number of intermediates populated by the kinase indicates that majority of the kinases undergo multistate folding. (f, g, h) Free energy (FE) profiles of a representative kinase from every family which are PRKACA from AGC, PIM1 from CAMK, CSNK1A1 from CK1, MAPK1 from CMGC, NEK1 from NEK, ULK3 from ‘Other’ family, MAP4K1 from STE, RIPK2 from TKL and ABL1 from TYR family. The uniqueness of these FE profiles indicates the presence of different interaction networks within each kinase that in turn determines the eventual folding-conformational landscape. Grouping of the family is done for better visualization.

The number of residues in kinases ranges between 240-330 (Figure 2b) and hence total number of microstates in the bWSME model range between 2-13 million (Figure 2c). We employ this large ensemble to predict the free-energy profiles of 274 kinases from the pruned database (see Methods and Figure 1c), as a function of the reaction coordinate (RC), the fraction of structured blocks (normalized to the total number of structured blocks in a given kinase). As a first step, we modulate the van der Waals interaction energy per native contact (ξ) to arrive at an overall stability of 30 kJ mol^-1^ for every kinase – this value is effectively the difference between the free-energies of the unfolded state (typically at RC values < 0.2) and the folded state (typically at RC values > 0.8) – and by keeping every other parameter constant other than the input structure. The values of ξ cluster between -70 to -85 J mol^-1^ per native contact (Figure 2d), with an average value of -76.5 J mol^-1^.

The free-energy profiles are always multi-state (Figure 2e, S2) with several partially structured states evident in the one-dimensional representation (Figure 2f-2h). A collection of shallow wells is observable between the RC values of 0.2 and 0.5, highlighting a stepwise folding mechanism wherein a specific region of the kinase folds first following which the rest of the structure assembles. The depth of the minima, however, varies depending on the kinase. The folded-like conformational substates (at RC>0.6) also show a large diversity: a small thermodynamic barrier separates two minima in PRKACA and PIM1 (Figure 2f at RC > 0.8), a large thermodynamic barrier separates the minima in MAPK1 (Figure 2g at RC > 0.6), or multiple sub-states co-exist with equal probabilities in RIPK2 (Figure 2h at RC > 0.8). The folding thermodynamic barrier can be as large as 30 kJ mol^-1^, but with multiple intermediate states which act as kinetic traps (Figure 2e).

### Local stabilities are anisotropically distributed

Given that the free-energy profiles are diverse, we next ask if the 274 kinase structures exhibit any patterns in their local stabilities. To estimate this quantity, we quantify the mean stability, ⟨Δ*G*_*s*_⟩, of N- and C-terminal lobes (see Methods) from the residue (block) folding probabilities. The N-terminal lobe (N_t_) displays a broad distribution of stability values ranging from stable (negative values) to unstable (positive values; Figure 3a). On the other hand, the C-terminal lobe (C_t_) is predominantly more stable with a sharp distribution of stabilities. The difference in stability is also observed at the level of individual families indicating a robust pattern (Figure 3b). The observed difference is a consequence of nearly 25% stronger packing density within the C_t_ compared to the N_t_ (Figure 3c, S3). In addition, the overall electrostatic interaction energy between the charged residues in the C_t_ is stronger than the N_t_ – this could be either the cause of the strong packing or the outcome and cannot be further deconvoluted (Figure 3d, S3).

**Figure 3.**
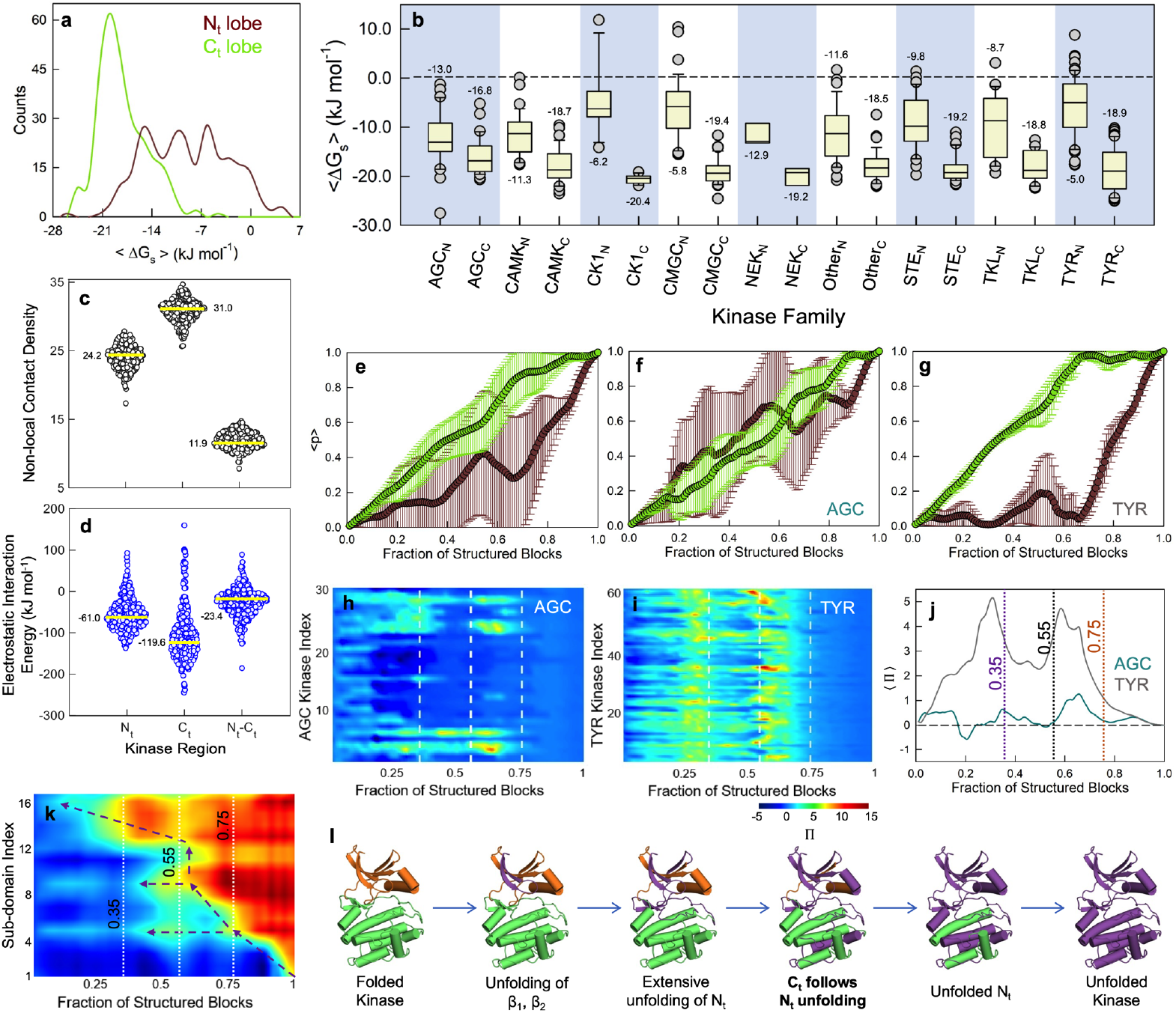
Anisotropic distribution of local stabilities. (a) Distribution of average lobe stability (⟨*ΔG*_*S*_⟩). The more negative mean value and the narrow spread of the C_t_ lobe indicates that it is more stable than the N_t_ lobe. (b) Box plot of ⟨*ΔG*_*S*_⟩ for the N_t_ (subscript ‘N’) and C_t_ lobe (subscript ‘C’). The median line (median value is mentioned) is included in the box plot, where the box denotes the interquartile range (IQR) with quartile 1 as lower and quartile 3 as the upper limit. The outliers denoted by circles are the data lying outside 10-90% of the population range denoted by the whisker caps. (c, d) A swarm chart of the non-local contact density (panel c) and electrostatic interaction energy (panel d). The median line is in yellow. (e, f, g) The mean folding probability graph of residues in the C_t_ and N_t_ lobes shows that the C_t_ lobe folds earlier (panel e). The error bars indicate the spread from all kinases. In the AGC kinase (panel f), the residues in N_t_ and C_t_ lobes fold concurrently. In contrast, the residues in C_t_ lobe of TYR kinase family (panel g) initiates folding prior to the residues of N_t_ lobe. (h, i) Plot of log ratio of folding probability of C_t_ and N_t_ lobe at each reaction coordinate (*Π* = *ln*(*p*_*ct*_/*p*_*Nt*_)) for each family member colored according to the color scale shown below. A value of zero (cyan-blue patch) indicates concurrent folding of both the lobes, while red or blue is representative of biased folding pathways dominated by N_t_ first (dark blue) and C_t_ first (dark red), respectively. The white vertical lines correspond to the specific points along the reaction coordinate discussed in the text. In AGC family (panel h), more blue-cyan patches are observed indicating concomitant folding, whereas more greenish-yellow patches are observed in TYR family (panel i) indicating C_t_-lobe folding first. (j) Mean of *Π* across all members (⟨*Π*⟩) of the AGC and TYR families from panels h and i, respectively, with specific reaction coordinate values highlighted. (k) The mean folding stability (⟨Δ*G*_*S*_⟩) of the 17 conserved kinase subdomains for all the kinases across the reaction coordinate. The dashed arrows represent potential unfolding pathways. A spectral color code is employed where red and blue represent the high and low stability, respectively. (l) A cartoon representation of the most-probable unfolding mechanism observed in kinases.

The differences in relative stabilities manifest as a generalizable folding mechanism wherein the residues in the C-terminal lobe folds first followed by the N-terminal lobe along the reaction coordinate, albeit with variations (Figure 3e, S4). Specific deviations are discussed here from the perspective of the two extreme cases: AGC and TYR kinase family members. In the AGC family, the domain folding probabilities overlap significantly along the folding coordinate, highlighting the coupled nature of the folding process or concomitant folding (Figure 3f). On the other hand, the TYR kinase family members almost always fold in a manner wherein the C-terminal lobe folds first to act as a scaffold for the N-terminal lobe to fold (Figure 3g). The trends are maintained at the level of individual members of these two families, and this can be observed in the plot of log ratio of folding probability of C_t_ and N_t_ lobe at each reaction coordinate defined by the variable (*ΠΠ* = *ln*(*p*_*Ct*_/*p*_*Nt*_)) for every family member (Table S3). In AGC family (Figure 3h), *Π* ≈ 0 across the reaction coordinate (blue-cyan patches) indicating concomitant folding, whereas *Π* > 1 in TYR family (Figure 3i) and is a consequence of C_t_-lobe folding first; the corresponding averages highlight the stark differences in the folding mechanism (Figure 3j). In fact, only AGC and CAMK family members exhibit this coupled folding mechanism, while the rest of the 7 families fold predominantly in the biased C_t_-first folding mechanism (Figure S4-S6).

The states sampled at different lower values of the RC provide a view of the weakly structured regions in the kinase; importantly, such regions which have been shown to be functional relevant across multiple protein systems.^41^ At a RC-value of 0.75, the folding probability averaged across all the 274 kinases indicates partial unfolding at the N_t_-lobe (Figure 3k, 3l), with the first four sub-domains harboring β_1_, β_2_ and β_3_ sampling partially structured states. These sub-domains are located at the interface between the two lobes and is the precise structural location where the cofactors (Mg^2+^, ATP) and substrate bind. At even lower values of RC (0.55), the sub-domains αC, β4, β5 and αD exhibit higher unfolding probability, followed by progressive unfolding of the C_t_-lobe (RC <0.35). Taken together, though the free energy profiles indicate varied partially structured states, there are generalizable patterns in local stabilities and hence (un)folding mechanism across the 274 protein kinases.

### Activation modulates the stability across the entire structure and not just locally

Kinase activation involves the formation of an unbroken C-spine region mediated by ATP and the structural re-arrangements involving G-loop, activation loop and the αC helix, which enables catalysis (Figure 4a, 4b). The closer packing of the spine residues is expected to enhance the stability of the active kinase conformations. As a representative example, we discuss PRKACA kinase from the AGC family. The active kinase conformation of PRKACA exhibits a higher gradient towards the folded state (green in Figure 4c, S7) relative to the inactive state. The regions corresponding to the C-spine, the phosphate-binding loop (G-loop), and the associated beta-sheets (sub-domains 1-3) are partially unstructured and appears as an intermediate on the reaction coordinate (Figure 4d). The presence of ATP ‘completes’ the C-spine, folds the phosphate-binding loop and thus destabilizes the intermediate. However, the expected trend of active being more stable than the inactive is, interestingly, not observed across all family members, with only ∼57% being more stable in the active form (Figure 4e). This can also be seen in the comparison of mean local stabilities across the 104 active-inactive pairs studied, with most active and inactive kinases exhibiting similar values with a tail towards lower stability values only for a small subset of inactive kinases (⟨Δ*G*_*s*_⟩ > 0; Figure 4f). However, all the catalytically important structural elements – R-spine, C-spine, HRD and DFG motifs – exhibit a much smaller and surprisingly similar spread in local stabilities in the active state conformation, compared to the inactive state (Figure 4g).

**Figure 4.**
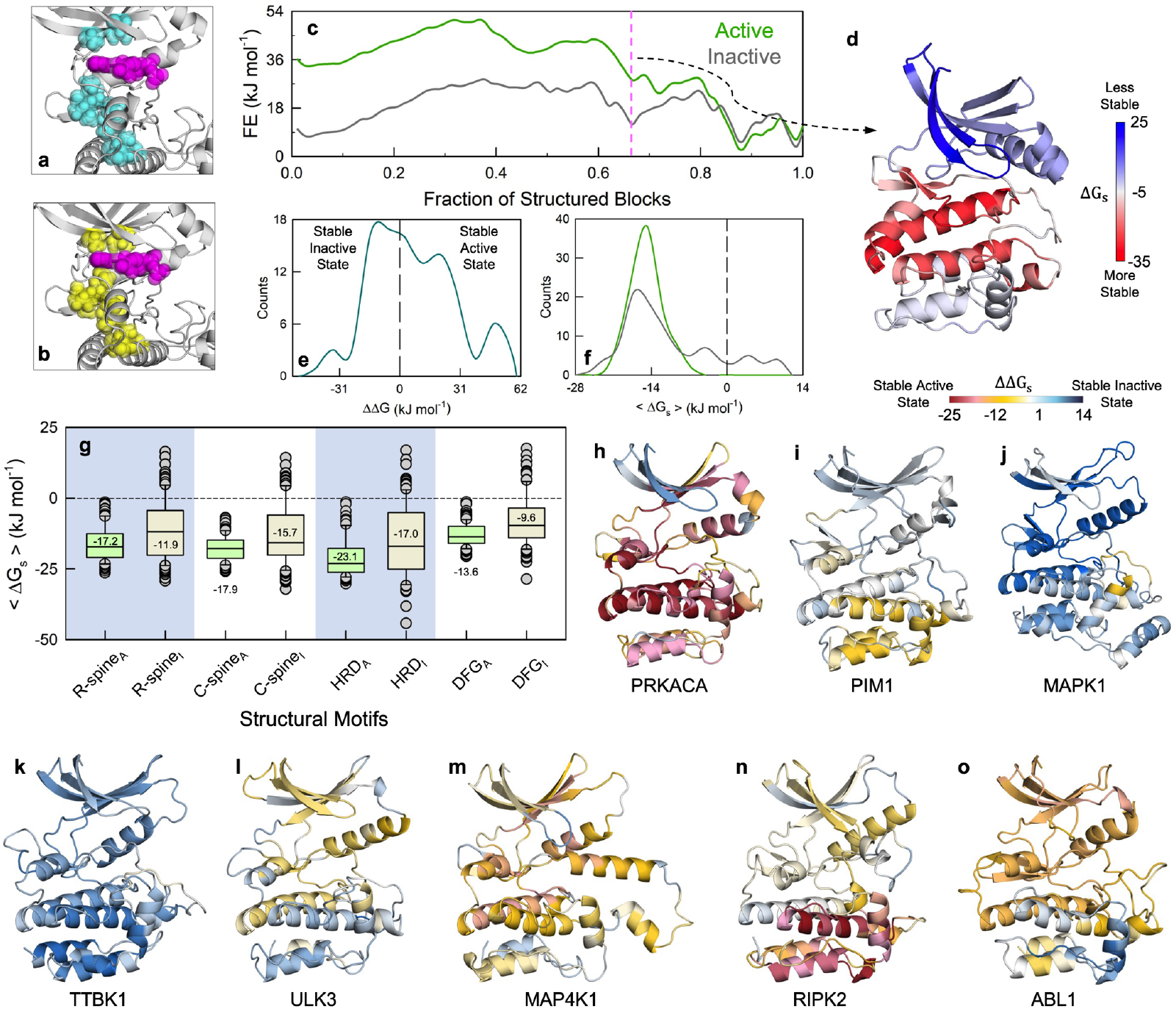
Changes in local stability patterns are not conserved between pairs of active-inactive kinases. (a) The broken C-spine residues (cyan + magenta spheres) of inactive state kinase structure (PDB id: 4AE6). The magenta spheres belong to adenosine triphosphate (ATP) atoms. (b) The active state kinase structure (PDB id: 3OVV) consisting of unbroken C-spine residues (yellow + magenta spheres). (c, d) Free energy profile of both active and inactive PRKACA (panel c). The reaction coordinate value (RC = 0.67) where an intermediate state is populated is marked by a pink vertical line. The corresponding structure at this RC-value is shown in panel d, with the local stability mapped. The blue region (primarily N_t_-lobe) is involved in the formation of C-spine, controlling the entry of ATP molecule. (e) Distribution of difference in global stabilities between active and inactive kinases. (f) Distribution of mean local stability (⟨Δ*G*_*S*_⟩) for all kinase residues across active (light green) and inactive (light grey) pairs. The narrow spread and more negative mean value suggest that the active state of majority of the kinase is more stable locally. (g) Mean local stability (⟨Δ*G*_*S*_⟩) of catalytically important motifs for active (subscript ‘A’, light green) and inactive kinases (subscript ‘I’, light grey). The median line is included in the box plot, where the box denotes the interquartile range (IQR) with quartile 1 as lower and quartile 3 as the upper limit. The outliers denoted by circles are the data lying outside 10-90% of the population range denoted by the whisker caps. (h-o) Difference in local stabilities (ΔΔ*G*_*S*_) between active and inactive kinase for eight representative kinases, one from each family There is no representative kinase from NEK and RGC family due to the absence of active-inactive structure pairs.

It is, however, likely that different kinase regions respond in a non-conserved manner to activation. In fact, the non-conserved response to activation can be vividly observed when the *differences* in local stabilities between active and inactive kinases, i.e. ΔΔ*G*_*s*_ = Δ*G*_*s,active*_ - Δ*G*_*s,inactive*_, are projected on to the structure; no two kinases are similar (Figure 4h-4o). The unique conformational preferences of the catalytically crucial motifs in the active state (and hence the similar stability values) therefore contribute to a redistribution of stabilities across the entire structure and not just in the immediate vicinity of the motifs, which is captured by the model. Many regions are indeed more stable in the active conformations (ΔΔ*G*_*s*_ < 0; red, orange or white in Figure 4h-4o). Thus, the inactive conformations are intrinsically more flexible, potentially primed for accepting co-factors and ligand and even phosphorylation. There are, however, exceptions. For example, in MAPK1 kinase, many regions display ΔΔ*G*_*s*_ > 0, indicative of more stability in inactive conformation than the active (Figure 4j). The MAPK1 kinase belongs to the CMGC family and majority of kinases in this family are regulated by other kinases,^43^ thus requiring a more detailed study.

### Allosteric Connectivity in Kinases

One of the standout features of kinases is the presence of multiple allosteric sites and mechanisms that regulate activity.^44–47^ The allosteric regions include sites for drug-binding (MT3, MEK1/2 type III inhibitor), activation (AAS, Aurora A activation segment), substrate binding (PDIG, recognition site near this PDIG motif), PDK1 regulation (PIF, PDK1 interacting fragment), myristoyl binding (CMP, c-Abl myristoyl pocket), auto-inhibition (MPP, MKK4 p38a peptide) and substrate recruitment (DRS, D-recruitment site present in MAPKs) (Figure 5a, S8).^48^ The allosteric sites are located on the solvent-exposed surface of kinases and interestingly located primarily in the N_t_-lobe. The mean local stabilities (⟨Δ*G*_*s*_⟩) of the allosteric sites are varied and spread over larger range in the inactive conformations compared to the active conformations (Figure 5b). Furthermore, not only are the medians marginally lower, but also ⟨Δ*G*_*s*_⟩ is tailed at higher values in the inactive conformations, indicative of weaker stability and by extension, larger dynamics. These observations mirror the behavior of the orthosteric site residues. Upon activation, it thus appears that the dynamics are constrained either through stronger packing or adopting an alternate conformation, and this feature is uniquely conserved across all kinases irrespective of their families.

**Figure 5.**
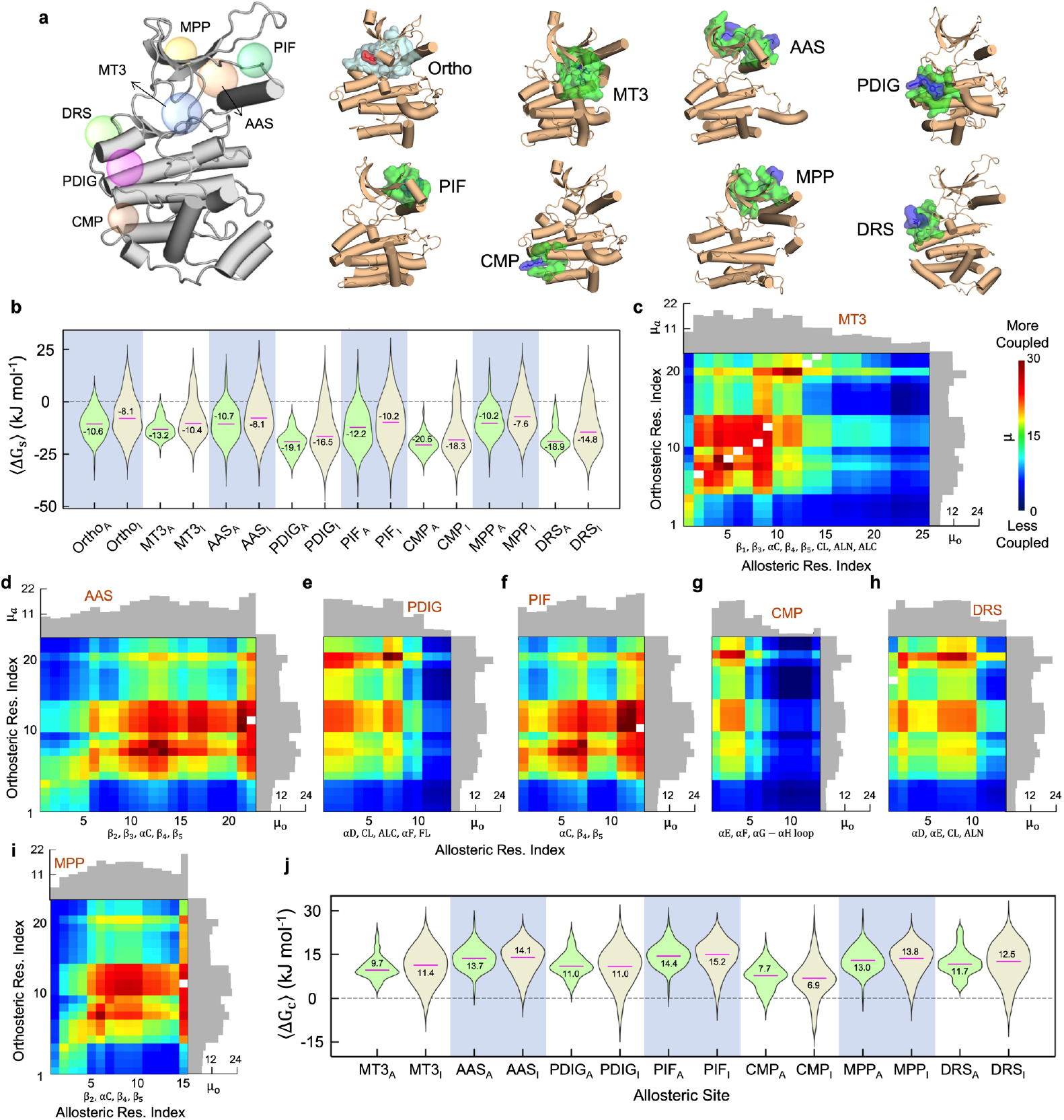
Allosteric connectivity in kinases. (a) Allosteric pockets in kinases together with the orthosteric site (cyan). (b) Mean local stability (⟨Δ*G*_*S*_⟩) of the different sites for the 104 active (subscript ‘A’, light green) and inactive kinase (subscript ‘I’, light grey) pairs. The spread of ⟨Δ*G*_*S*_⟩ for orthosteric and allosteric residues are broader in the inactive kinases compared to that of active kinases, suggesting stabilization of these sites upon activation. The median value of the distribution is in pink. (c-i) The two-dimensional colormap of mean effective thermodynamic coupling across 274 kinases (µ) for each allosteric site against the orthosteric site. The mean effect of all the orthosteric residues on each allosteric residue (µ_*a*_) is shown by histogram above the plot, while mean effect of all the allosteric residues on each orthosteric residue (µ_*o*_) is shown by histogram on the right. White squares represent residues that are common between both ortho- and allosteric sites. (j) Mean effective thermodynamic coupling free energies (⟨Δ*G*_*c*_⟩) of residues in the ortho and allosteric sites in both active (subscript ‘A’, light green) and inactive kinase (subscript ‘I’, light grey) pairs.

We next ask if there are trends in the extent of the coupling between pairs of residues – one located in the orthosteric site and the other in the allosteric site (Table S4). These pairwise effective thermodynamic coupling free-energies (Δ*G*_*c*_) are calculated from the ensemble of microstates and their relative folding probabilities thus providing unique insights into the underlying thermodynamic architecture that dictates allosteric connectivity; higher and lower pairwise Δ*G*_*c*_ values are indicative of sites that are strongly and weakly connected, respectively. Three observations stand out. First, all the allosteric and orthosteric site residues are strongly coupled with each other, irrespective of their location. The corollary is that any perturbation to one residue in either of the sites will naturally be ‘felt’ at the other site (sea of red, green and cyan in Figure 5c-5i, S9). Second, the mean effect of the orthosteric residues on the allosteric site residues are varied (the distribution on top in Figure 5c-5i), as the identity of the allosteric site residues are different, which in turn leads to the non-conserved modulation. Third, a unique band of residues always light up (dark red in Figure 5c-5i) which span residues 6-14 and 20 in the orthosteric site index. The residues 6-14 include the conserved Lys from β_3_ which is acknowledged as a critical residue for signal propagation between ATP-binding pocket and an allosteric site,^49^ conserved Glu from αC that induces the kinase into active conformation, along with the other hydrophobic residues that stabilize ATP in the pocket. Residue 20 is a part of the conserved hydrophobic C-spine (Leu in the majority of kinases^27^) whose orientation determines whether a kinase is catalytically primed.

The allosteric maps in panel 5c-5i are averaged across all the active-inactive pairs and plotted in Figure 5j. The mean effective thermodynamic coupling free energies now display a trend expected from ⟨Δ*G*_*s*_⟩: the thermodynamic coupling *between* orthosteric and allosteric sites spans a larger range of values across the different kinases in the inactive conformation compared to the active conformation. The tail of the distribution is also observable at lower values of Δ*G*_*c*_ in the inactive conformation, while the active conformations exhibit similar values of Δ*G*_*c*_ signifying similar structures (and side-chain orientations). On average, the connectivity between the orthosteric and allosteric sites are weaker in the inactive form compared to the active form, uncovering hidden differences in allosteric connectivity which is not obvious when the structures are superimposed.

### Diverse intra-molecular interaction networks contribute to the varied conformational behavior

The rich conformational behavior of kinases evidenced by the dissimilar free-energy profiles, and varied extents to which different parts of structure are perturbed upon activation are naturally a consequence of the diverse intra-molecular interaction network. In other words, apart from generic differences in the stability of the two lobes, small differences in sequences distributed across the structure can manifest as large changes in connectivity patterns. One avenue to test for this is to extract differential coupling indices (DCIs) by mutating a residue to alanine and plotting the effect of this mutation on the overall landscape, i.e. an alanine scan. From the perspective of the model, this is done by quantifying the difference in positive coupling free energy between the WT and the mutated variant (|ΔΔ*G*_+_|, see Methods and Figure 6a). The positive coupling free energy carries information on the pair-wise thermodynamic coupling between different residues that are simultaneously folded across the millions of microstates; the folded status is in turn governed by the nature and strength of interactions mediated by the constituent residues both from direct (first shell around a residue) and indirect (second shell and longer) interactions.

**Figure 6.**
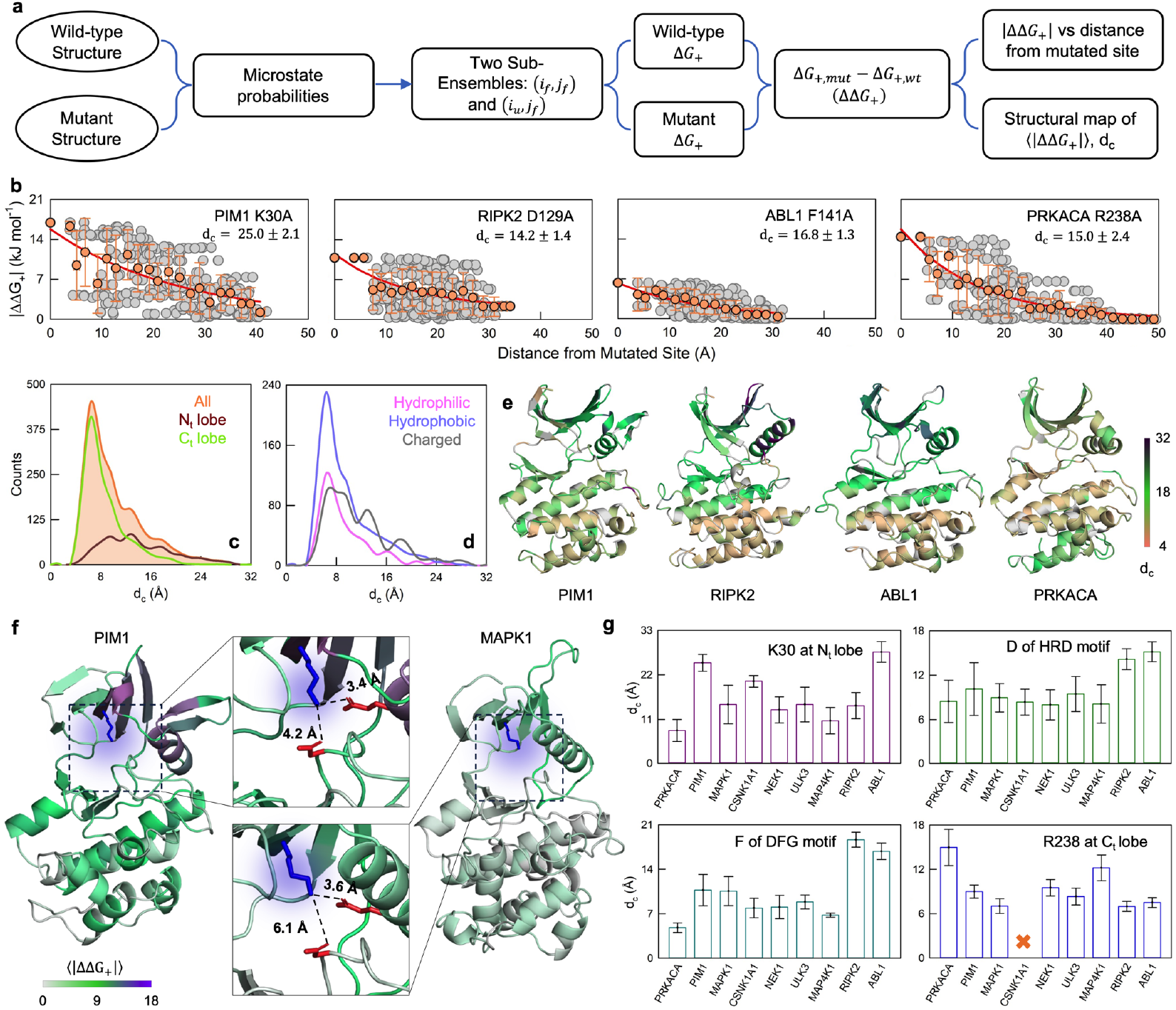
Pervasive long-range thermodynamic coupling in kinases. (a) Workflow of the methodology utilized for the alanine scanning mutagenesis. (b) Absolute mutational response (⟨|ΔΔ*G*_*i*_|⟩) with respect to distance from mutated site for four representative mutational sites selected from the N_t_ lobe, catalytic loop, activation loop and C_t_ lobe. (c) Distribution of coupling distances (*d*_*c*_) for all the mutated residues (orange), residues of N_t_ lobe (brown) and residues of C_t_ lobe (green). (d) The distribution of *d*_*c*_ for hydrophilic (pink), hydrophobic (blue) and charged residues (gray). (e) The magnitude of coupling distance is mapped onto the structure for the kinases shown. Residues in the N_t_ are more coupled to distant residues (higher *d*_*c*_) than the residues in C_t_. (f) Structural map of ⟨|ΔΔ*G*_*i*_|⟩ for wildtype PIM1 (PDB id: 1XWS) and wildtype MAPK1 (PDB id: 3SA0). The mutated residue is shown as a blue, glowing stick while the negatively charged residues in proximity are shown as red sticks. The color ranges from grey to violet, where residue with maximum mutational effect is colored violet, whereas regions with nearly zero mutational response are colored as grey (color bar below). (g) Bar graph of the coupling distances (*d*_*c*_) at structurally identical position for nine kinases (except RGC due to the unavailability of experimental structures). The structural positions correspond to K30, D124, F143, and R238, respectively, in PRKACA.

Representative examples of DCIs for specific residues are shown in Figure 6b (see Figures S10-S17 for the full set). Note that when the |ΔΔ*G*_+_| is binned at every 2 Å, the resulting mean follows an exponential-like trend which is the zeroth order effect expected upon perturbations, as observed across diverse experiments and model systems.^50,51^ Such DCIs were generated for nine kinases (one member from every family) amounting to 1975 mutants, and the distance dependence was fitted to exponential functions to extract the coupling distance, *d*_*c*_. The resulting coupling distances vary from 2-28 Å with a heavy right-tailed distribution. The *d*_*c*_ distributions display a peak value at 7 and 12 Å, respectively, for C_t_- and N_t_-lobes, highlighting that the effect of any perturbation in the N_t_-lobe which harbors most of the activity-determining residues is felt to a longer distance compared to the C_t_-lobe (Figure 6c, Figure S18-S22). This specific observation potentially explains why most allosteric pockets are in the Nt-lobe. Mutations involving both hydrophobic and charged residues have a larger effect on the ensemble as intuitively expected and as observed earlier in smaller domains (Figure 6d, Figure S18-S22). Thus, the distribution of *d*_*c*_ follows a pattern similar to trends in stability, on average, but with differences across the kinase members from every family (Figure 6e, Figure S23).

The question is then: how different are the interaction networks within every kinase? We answer this by quantifying the mutational response of structurally and catalytically important residues located at identical structural positions. If the packing density and interactions are similar, the effect of perturbation is expected to be similar for a given position across kinases. We consider four residues - K30 in the Nt-lobe, D in HRD motif, F in DFG motif and R238 in the C_t_-lobe of PRKACA (AGC family) – for further discussion. In PIM1, the K30A mutation results in a *d*_*c*_ of 25 Å, while the same K30A mutation in MAPK1 displays a *d*_*c*_ of just 14 Å. There is a shared salt-bridge between K30 and E52 and D149 in PIM1, which in turn brings together the loops in proximity to each other enhancing packing of nearby residues. In MAPK1, the salt bridges of K30 with E47 and D143 are present but separated by longer distances, and consequently the loops are weakly packed, as can be visually observed in Figure 6f. The mutation to alanine not only eliminates the salt-bridge but also the packing interaction, and the effective change is therefore more significant in PIM1 than MAPK1, contributing to a larger coupling distance and higher amplitude of perturbation in the former (Figure S24). Small changes in packing density can thus result in significant changes in positive thermodynamic coupling and this is observed across the 4 positions studied with the *d*_*c*_ varying anywhere between 5 – 25 Å (Figure 6g), apart from changes in amplitude (Figure S24). These results highlight the large differences in the nature and strength of interactions between different residues in kinases, which are not obvious in structural superimpositions or evolutionary studies, and the bWSME model provides a simple avenue to extract them.

### Abl1 kinase as a model system

As a case study, we predict the conformational features for a well-studied model system, the Abl1 kinase. NMR-structures of three of the functional forms of the kinase are available and they correspond to the active, inactive 1 (I1) and inactive 2 (I2) conformations, indicative of a complex native ensemble.^52^ Structurally, there are only minimal differences between the active and I1, while I2 exhibits a much larger change in the positions of the αC helix and the activation loop, rendering the kinase inactive.

The folding free-energy profiles (FEPs) exhibit a unique trend wherein the fully folded state (marked by a*) is the most populated in the active conformation followed by I2 and then I1 (Figure 7a). The FEPs of active and I1 are similar, but that of I2 differs markedly. The conformation b* with partial structure in the phosphate-binding loop is the ground state in I1, and this is in agreement with the lower stability expected for the N_t_-lobe. The FEP derived from I2 conformation is quite flat with the conformational substates b*, c*, and d* exhibiting similar free-energy values. The substates identified to the left of b* on the RC are progressively more unfolded at the Nt-lobe and adjacent regions. We thus predict that the kinase is highly flexible in the Nt-lobe regions where the ATP and substrate binds (I2 free-energy profile), but upon activation many such states are destabilized and the folded-like compact state is the most preferred (active and/or I1 free-energy profile; note the upward arrow). CEST-NMR experiments point to a large fraction of residues in the N_t_-lobe undergoing conformational exchange,^52^ confirming that the model can *de novo* identify partially structured states given a structure.

**Figure 7.**
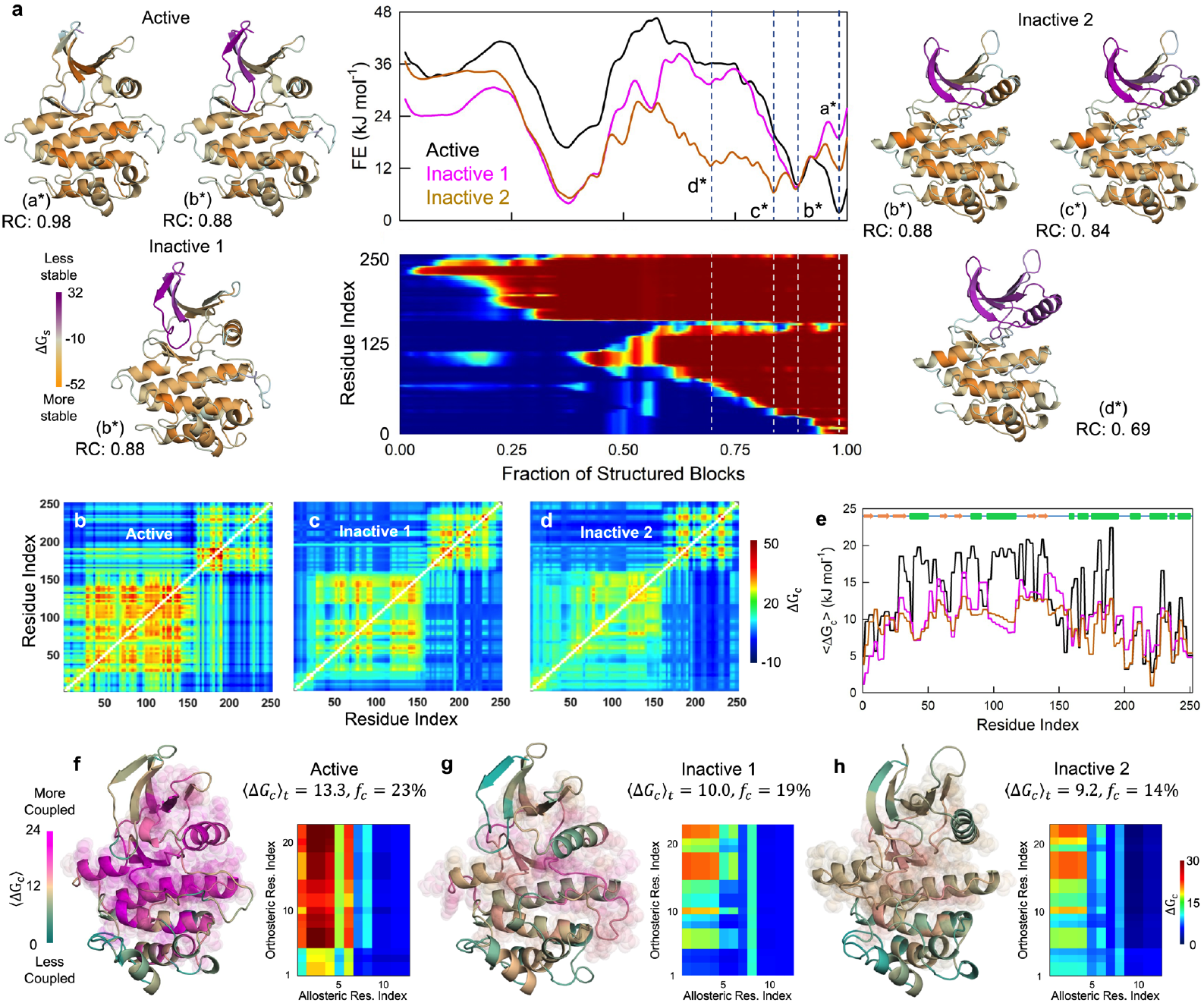
Multi-state conformational landscape of Abl1 kinase. (a) Free energy profiles of Abl1 kinase calculated from experimentally solved active, inactive 1 (I1), and inactive 2 (I2) conformations (top rectangular box). The structural features of the partially structured/intermediate states populated are depicted on either side of the free-energy profile. The active state primarily populates a* and to a minor extent b* (appears as an excited state), while the I1 state primarily populates b* with a* appearing as an excited state. The I2 state populates all the four partially structured states as the predicted free energy profile is flat at RC-values > 0.7. The structures of active (PDB id: 6XR6), I1 (PDB id: 6XR7), and I2 (PDB id: 6XRG) are colored based on the stability of the residues that ranges between orange (the most stable) to purple (the least stable). The two-dimensional plot below the free-energy profile represents the loss structure as a function of RC for the active state. A spectral color code from blue (unfolded) to red (folded) is employed. (b-d) Pair-wise effective thermodynamic coupling free energies (Δ*G*_*c*_) calculated from the different starting conformations. The strength of Δ*G*_*c*_ decreases from active to the completely inactive state. (e) The mean effective coupling (⟨Δ*G*_*c*_⟩) across the rows at each residue for all 3 states. The secondary structure elements are provided at the top of the plot with green box, salmon arrow and blue line representing helices, β-sheets, and loops. (f-h) Structures of the three Abl1 states, colored according to the ⟨Δ*G*_*c*_⟩, with residues having Z-scored Δ*G*_*c*_ > 1 are shown as colored spheres. The extent of thermodynamic coupling between orthosteric residues and the allosteric myristoyl binding site (CMP site) is shown in the lower right of every structure as a colormap. The spectral color code here ranges from 0 (dark blue) to 30 (dark red) kJ mol^-1^.

Studies on kinases have highlighted the role of synchronized dynamics across multiple protein regions in regulating activity.^53^ Does Abl1 kinase exhibit a similar feature? The pairwise coupling free energy matrices generated from each of the three starting conformations lead to diverse outcomes, with the active conformation being the most coupled followed by I1 and I2, both in the coupling free-energy matrices (Figure 7b-7d) and in the mean per-residue plots (Figure 7e). The strongly coupled residues of the active conformation (∼18-22 kJ mol^-1^) span the entire N_t_-lobe and a large fraction of the C_t_-lobe adjacent to the active site, which is the cleft between the two lobes. However, the inactive 2 conformation derived coupling free energies are marginal ranging from 6-12 kJ mol^-1^ (relative to that of the active conformation). In fact, the fraction of strongly coupled residues (*f*_*c*_), quantified by a Z-scored Δ*G*_*c*_ > 1 constitute 23% of the total residues in the active conformation but decreases to 14% in the I2 conformation (Figure 7f-7h). The *f*_*c*_ values of the I2 are within the range expected for proteins that range from 8-24% with a mean of 16%.^41^ However, the *f*_*c*_ in the active conformation is on the higher side with only CI2 (22%) and Kemp-eliminase (24%) matching the numbers,^41^ indicating a unique structural feature not observed in most proteins. Despite the graded change in coupling patterns, the allosteric myristoyl binding site^54^ is still thermodynamically coupled to the orthosteric site (the coupling maps at the bottom right of every structure in Figure 7f-7h), and follows the trends reported in Figure 5.

## Conclusions

In this work, we address the question of the extent to which small sequence changes modulate conformational landscapes by taking the challenging case of eukaryotic protein kinases. We employ the detailed bWSME model of protein folding to predict the folding conformational landscapes, local stabilities, coupling free energies and quantify the sensitivity to perturbations. The sequence diversity of kinases leads to diverse conformational behavior across the family members with different populations of partially structured states, arising from differences in the intra-molecular interaction network (Figure 2). Despite this, we find that the unfolding mechanism of kinases is conserved with the N-terminal lobe unfolding first followed by the C-terminal lobe (Figure 3), and this is a consequence of the anisotropic distribution of stabilization free energies across the two sub-domains. The populated partially structured states in the native side of the folding barrier exhibit different degrees of structuring in the N-terminal lobe, with the partially folded phosphate-binding loop (G-loop) appearing as an intermediate consistently across the kinase family. This feature is expected to modulate the dimensions of the cleft (where ATP and Mg^2+^ bind) and is consistent with expectations from nanopore tweezer experiments on Abl1 kinase,^55^ HDX-MS experiments on PDK1,^49^ NMR-derived observations on the dynamics and importance of αC-β4 loop,^56^ and broadly align with earlier predictions on how functionally relevant regions are partially folded in the native ensemble leading to large dynamics which aid in ‘accommodating’ the ligand.^41,57^ Statistical coupling analysis of kinase sequences reveal that residues in the cleft act as one continuous co-evolving unit.^58^ Taken together with our results, it is likely that mutations (specifically mutations associated with cancer^58^) in the cleft could not only modulate the binding strengths to co-factors/substrate but also tune the open-close equilibrium, which might be critical for selectivity and fine-tuning of activity.

In kinases, as with GPCRs,^38^ specific functional motifs including phosphate binding/G-loop, hinge loop, DFG motif and activation loop are all highly conserved and modular.^11^ On the other hand, the sequence changes at the other structural regions act in unison to modulate the output in a targeted manner, which in turn leads to diverse conformational substates in the native ensemble in the active or inactive forms that are reflected in the local stability patterns (Figure 4). Allostery requires an exquisite balance between the degree of conformation redistribution across these substates and the eventual functional output. This would require a fine-tuned intra-molecular interaction network that responds to various ambient cues and molecules. The highly coupled residues – quantified by the coupling distances (*i*.*e*. the extent to which the effect of mutations is felt) - in kinases cluster around the active site, i.e. at the N_t_-lobe and at the interface of the N_t_ and C_t_-lobes (Figure 5, 6), aligning with the known allosteric pockets. Such large coupling distances intrinsic to catalytically important structural regions might have enabled the evolution of allostery in such systems, as small sequence changes distance from the active site would intrinsically modulate the strength of the interaction network, and hence the connectivity, and finally functionality.

One of the defining features of the allosteric pockets in inactive kinases is their lower stability and thermodynamic coupling free energies, relative to the active kinases (Figure 5). Regions of lower coupling free energies generally exhibit enhanced conformational dynamics, and in kinases they are distributed primarily throughout the surface regions of the N_t_-lobe. These are not too dissimilar to the cryptic pockets that open-up transiently in proteins^59^ but indicate that it is crucial to map the native dynamics to identify structural regions that are less stable. These regions can then be targeted in a rational manner via docking small molecules. The myristoyl-binding pocket in Abl1 kinase is the only site that appears to be farther from the orthosteric site with an orthosteric-allosteric centroid distance of ∼25 Å. Despite the longer distance, myristoyl binding does propagate to the binding site, defining activity. In fact, the trends in coupling free energies follow the long-range exponential-like patterns (as a function of distance from the active site) expected of perturbations in protein systems^50,51^ and observed in large-scale mutagenesis studies on Src kinase.^60^ Our results further explain why attempts at swapping residues only at the binding site of imatinib in Src-kinase with that of Abl-kinase is unable to also swap the sensitivity proportionately,^61^ and the ‘distributed’ nature of mutations that confer resistance to ATP-competitive inhibitors;^62^ these are a consequence of a residues at distant sites regulating binding through subtle influence on dynamics, consistent with our observations.

In summary, our work provides the first detailed thermodynamic map of eukaryotic kinase family, highlighting both conserved and non-conserved aspects of the conformational landscape. We showcase how sequence changes within paralogs can have strong effects on the ensemble, underscoring the role of the global interaction network in fine-tuning activity. Accordingly, the strong connectivity of active residues to a large fraction of kinase residues including the C-terminal lobe, makes the system intrinsically sensitive to functional perturbations. Given that the bWSME model requires only the structure of the protein as an input, it should be possible to identify regions of proteins that are strongly coupled to the active site, quantify the coupling distance, and use these sites as hot-spots for identifying drug-like molecules.

## Abbreviations

bWSME: block Wako-Saitô-Muñoz-Eaton;
DCM: differential coupling matrix;
DCI: differential coupling indices;
RC: reaction coordinate

## Acknowledgements

The authors are grateful for the support from the early career Institute Research and Development Award (IRDA, IIT Madras) to A. N. N. The authors acknowledge the High-Performance Computing Environment (HPCE) facility, IIT Madras, for providing computational facilities.

## Data availability

All datasets generated during this study are available in the Github repository, https://github.com/AthiNaganathan/Kinase-Landscapes.

## Code availability

The data analysis codes and scripts employed in this study used MATLAB. The basic algorithm, code used for generating free energy profiles, landscapes, and coupling free energy matrices are available at https://github.com/AthiNaganathan/Kinase-Landscapes. Any scripts required for analysis are freely available on request by contacting the corresponding author.

