## Supporting Information for "Diverse Conformational Ensembles Define the Shared Folding-Allosteric Landscapes of Protein Kinases"

AUTHOR INFORMATION

**Corresponding Authors**

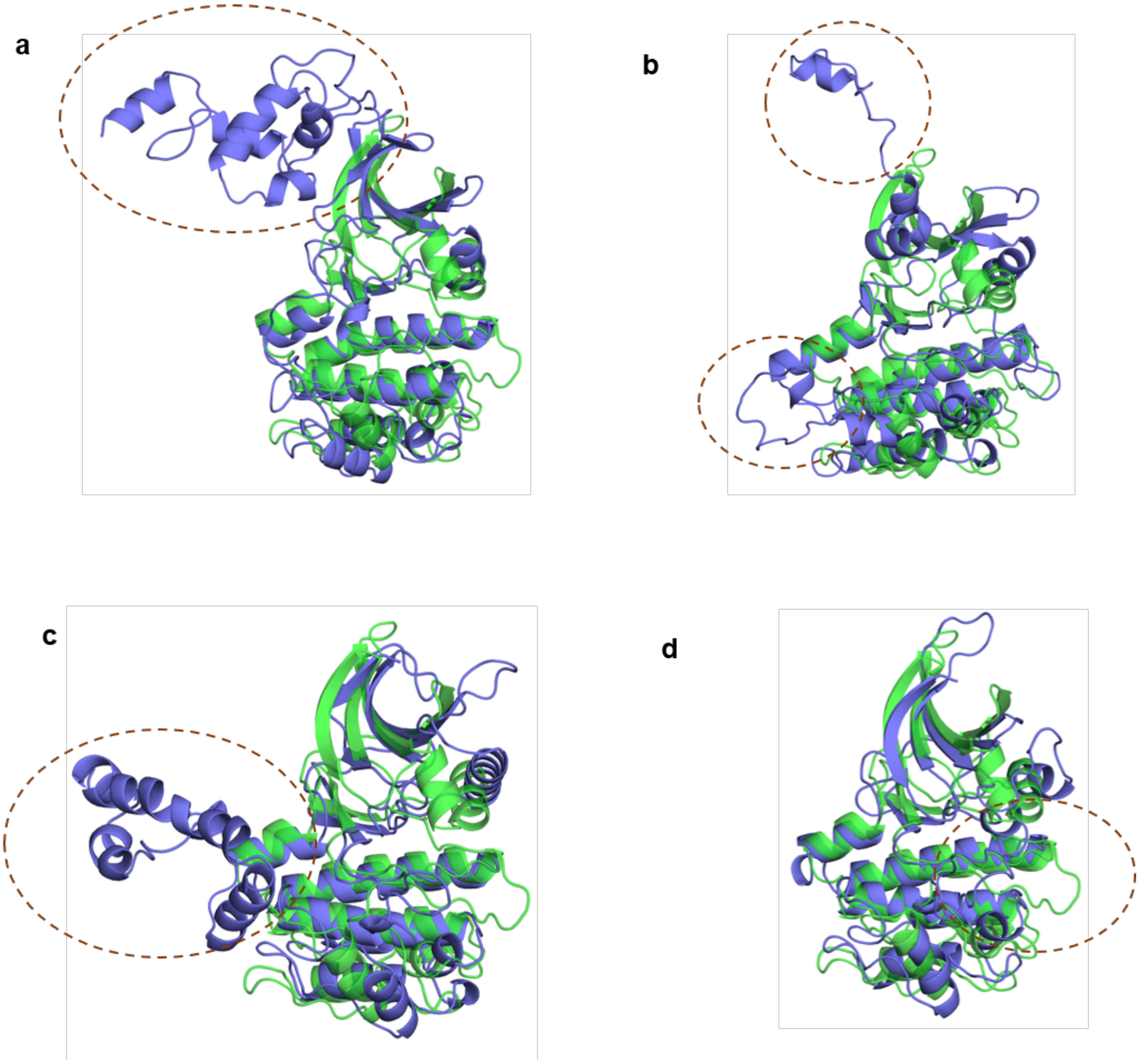


**Figure S1** Discarded eukaryotic protein kinases - TRIB1 (PDB id: 5CEM), PBK (PDB id: 5J0A), FLT1 (PDB id: 3HNG), ERBB3 (PDB id: 6OP9) - aligned with PRKACA (PDB id: 3OVV) in green. The unaligned regions are marked by brown dashed ovals. The structural alignment is performed with mTM-align.[1]

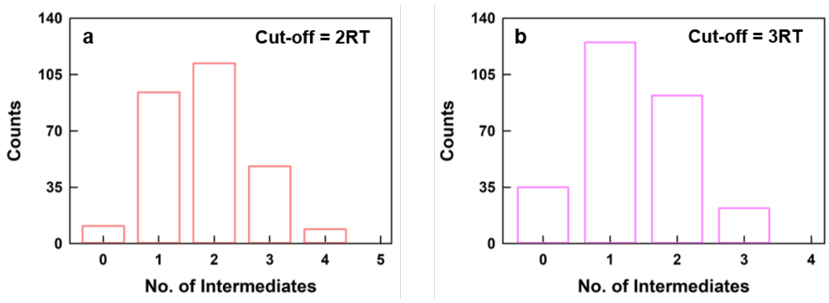


**Figure S2** (a, b) Bar plot for number of intermediates populated by the kinases when the valley depth cut-off is fixed to 2*RT* (panel a) or 3*RT* (panel b).

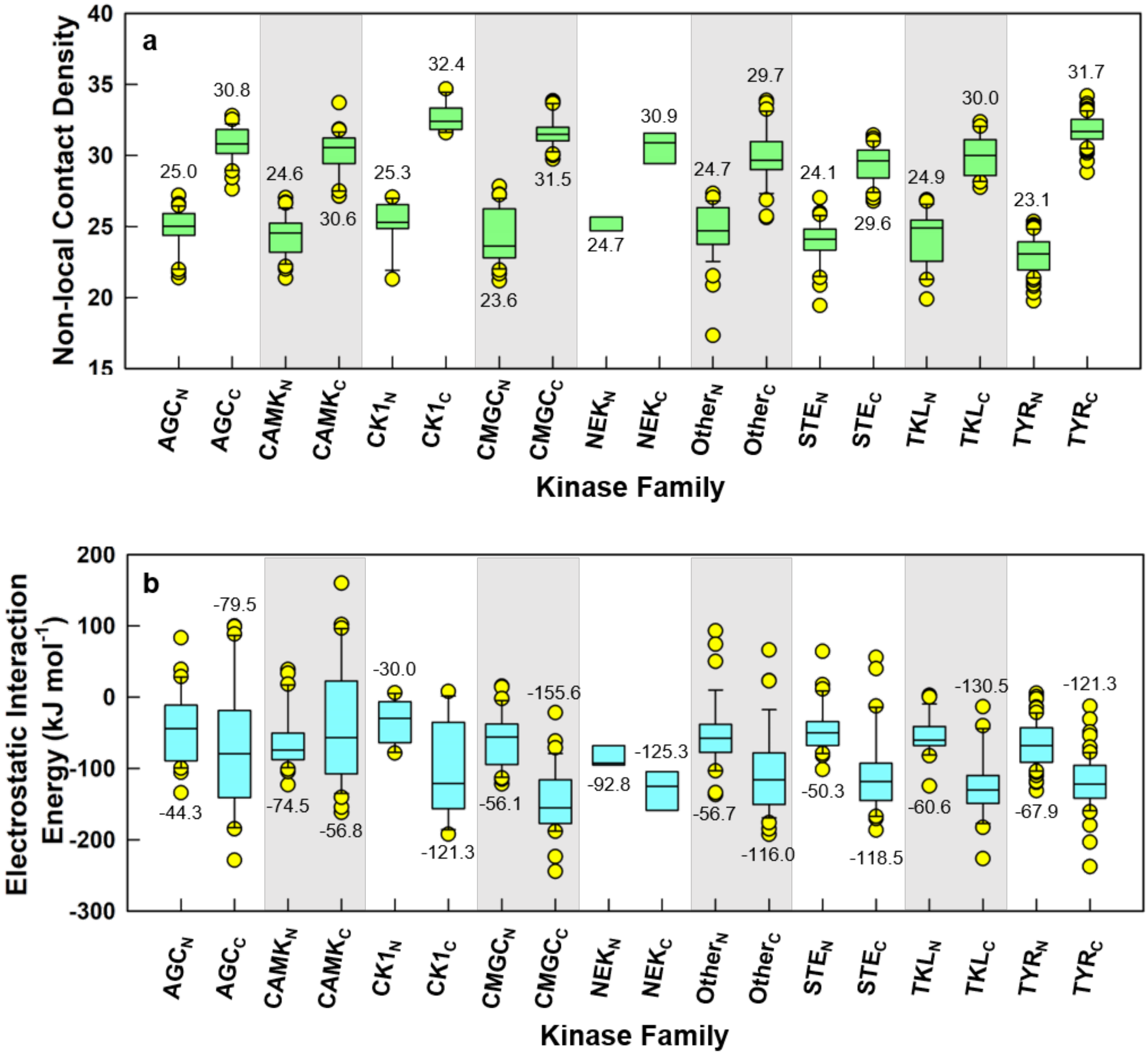


**Figure S3** (a, b) Non-local contact density (panel a) and electrostatic interaction energy (panel b) of $N_t$- (subscript 'N') and $C_t$-lobe (subscript 'C') for every kinase family. No experimental structures are available for the RGC family and hence is not used in this analysis. The median line (median value is mentioned) is included in the box plot where the box denotes the interquartile range (IQR) with quartile 1 as lower and quartile 3 as the upper limit. The outliers denoted by circles are the data lying outside 10-90% of the population range denoted by the whisker caps.

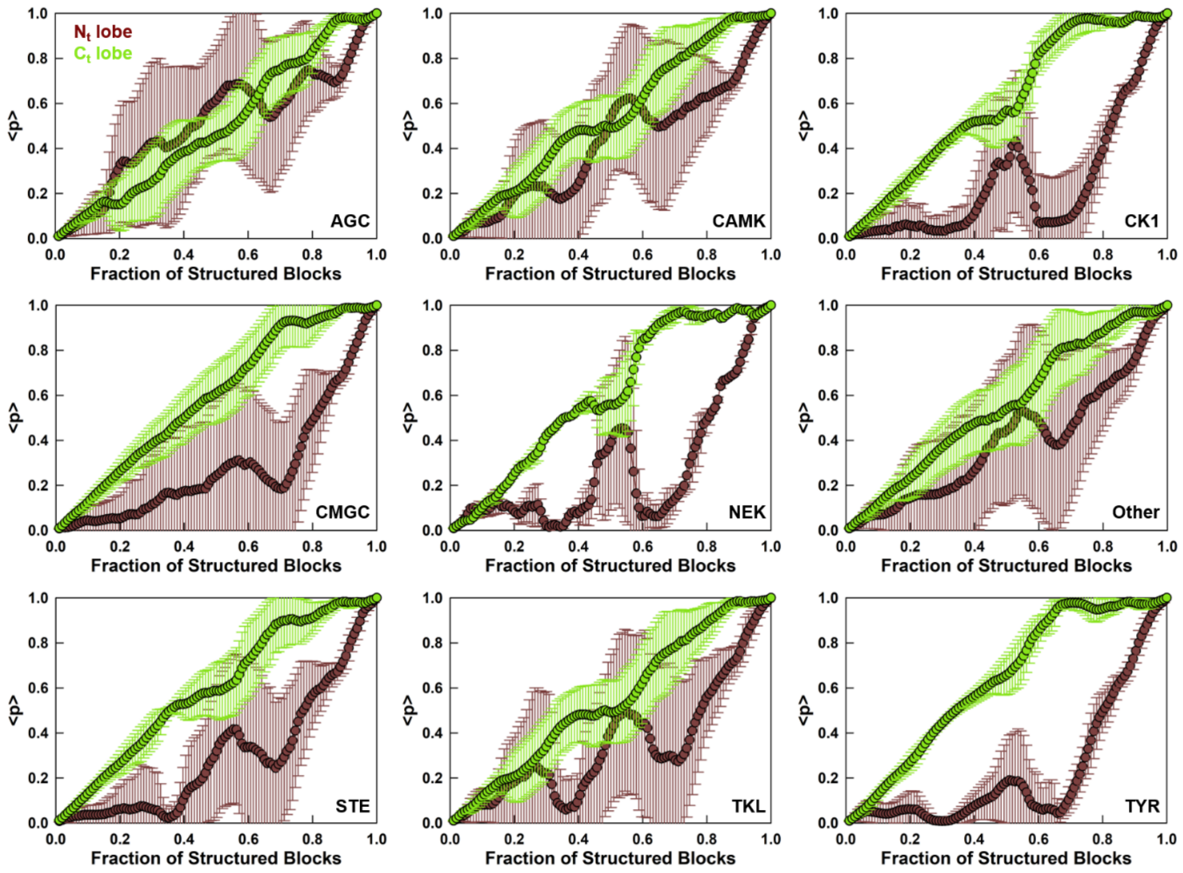


**Figure S4** Folding mechanism as predicted by bWSME model. The panels plot the mean folding probabilities ($\langle p \rangle$) of residues in the $N_t$- and $C_t$-lobe as a function of the reaction coordinate, for all the kinases except RGC kinase.

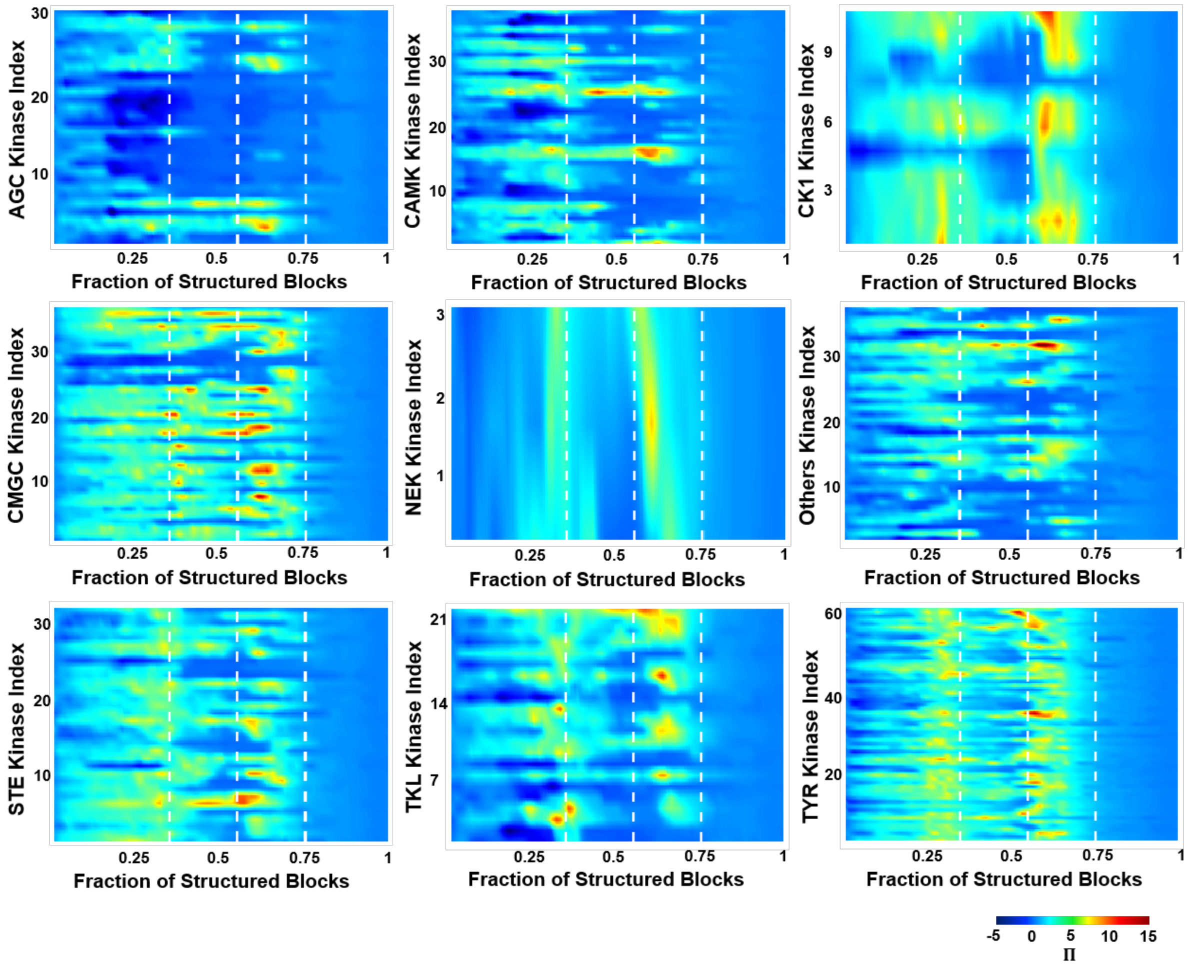


**Figure S5** Folding mechanism as predicted by bWSME model across all members of a given family. The plots are the log ratio of folding probability between $N_t$- and $C_t$- lobe residues $\left(\Pi = ln(p_{Ct}/p_{Nt})\right)$ as a function of the reaction coordinate, except for RGC kinase.

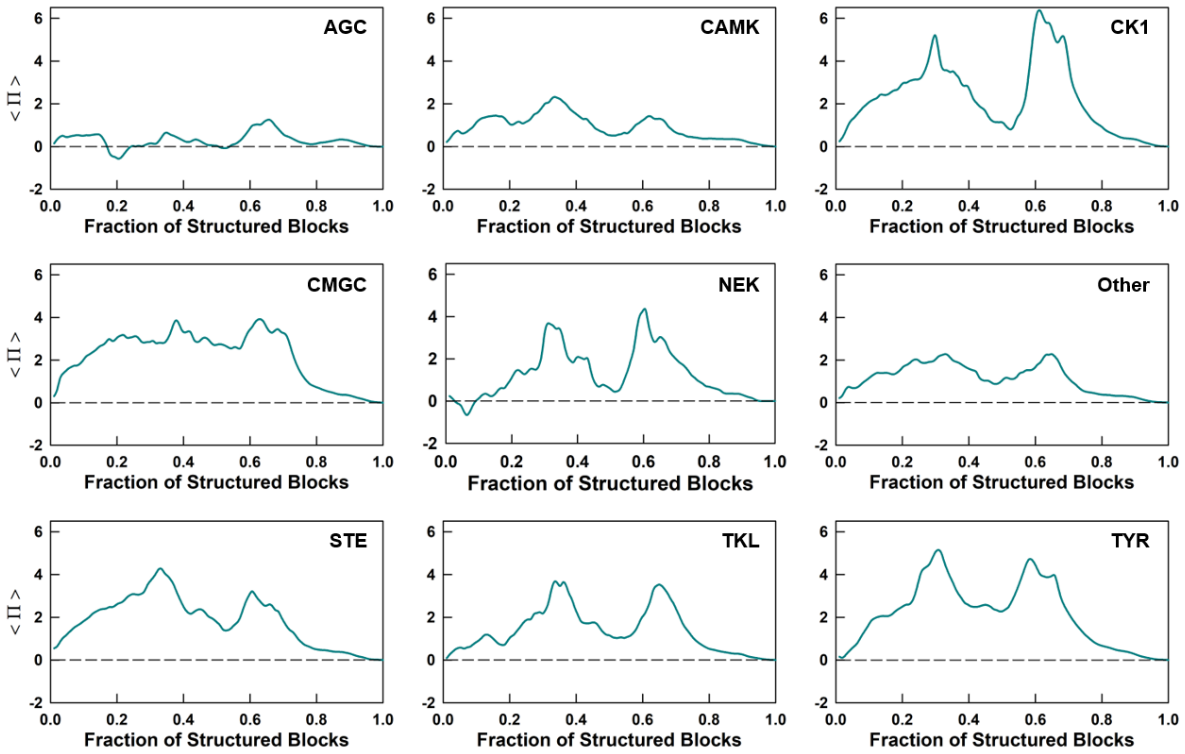


**Figure S6** Mean folding pattern as predicted by bWSME model. The *mean* of the log ratio of folding probability between $N_t$- and $C_t$-lobe residues ($\langle \Pi \rangle = \langle ln(p_{Ct}/p_{Nt}) \rangle$) as a function of the reaction coordinate is plotted, except for RGC kinase.

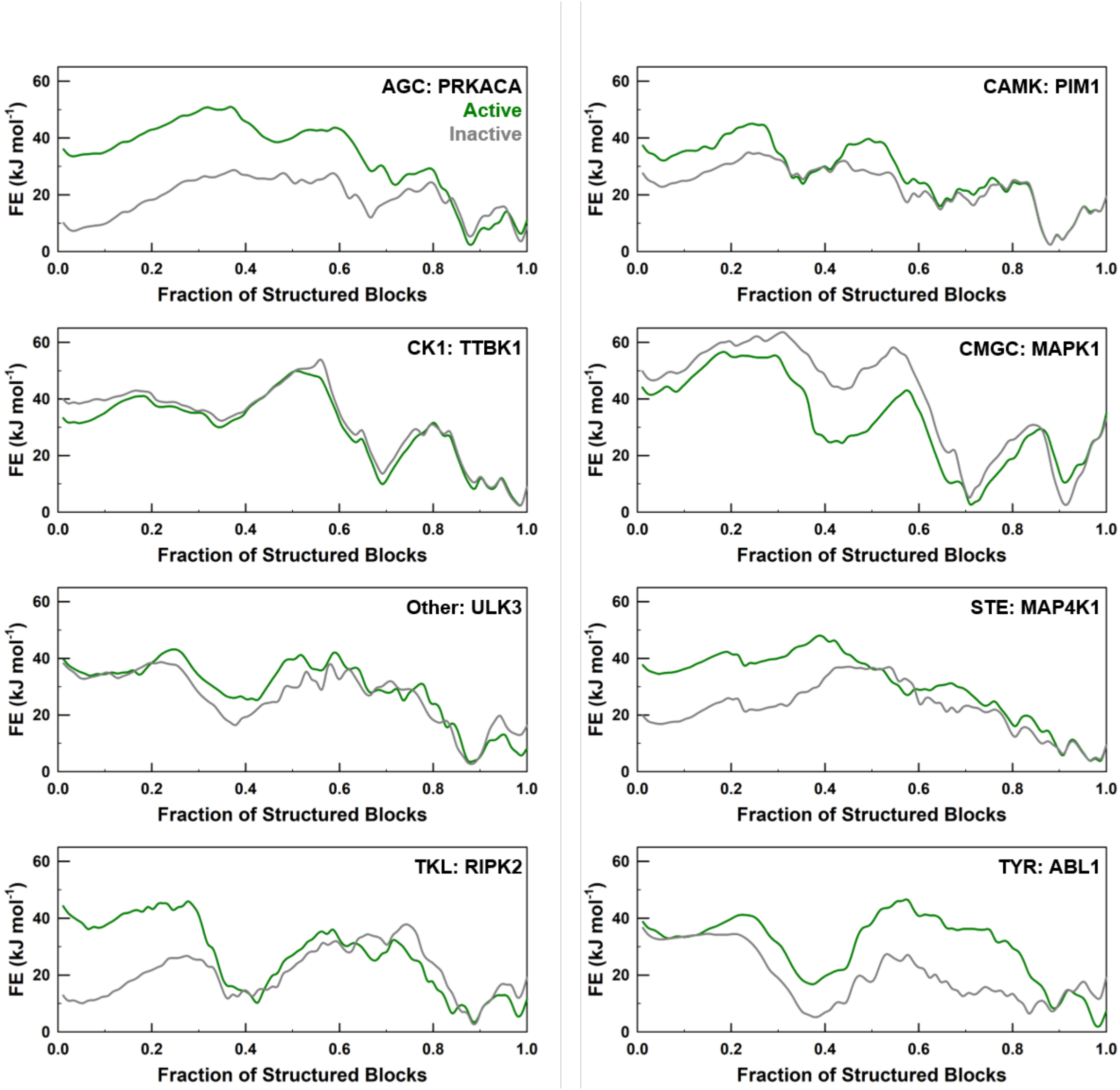


**Figure S7** Free energy profiles of representative active (green) - inactive (grey) kinases, one from each family. The PDB ids are: PRKACA - 3OVV (A), 4AE6(I); PIM1 - 1XWS(A), 6KZI(I); TTBK1 - 4NKN(A), 7Q8W(I); MAPK1 - 3SA0(A), 1WZY(I); ULK3 - 6FDY(A), 6FDZ(I); MAP4K1 - 7M0M(A), 7M0K(I); RIPK2 - 6S1F(A), 5NG3(I); ABL1 - 6XR6(A), 6XRG(I). Here ‘A’ and ‘I’ stand for active and inactive conformations, respectively.

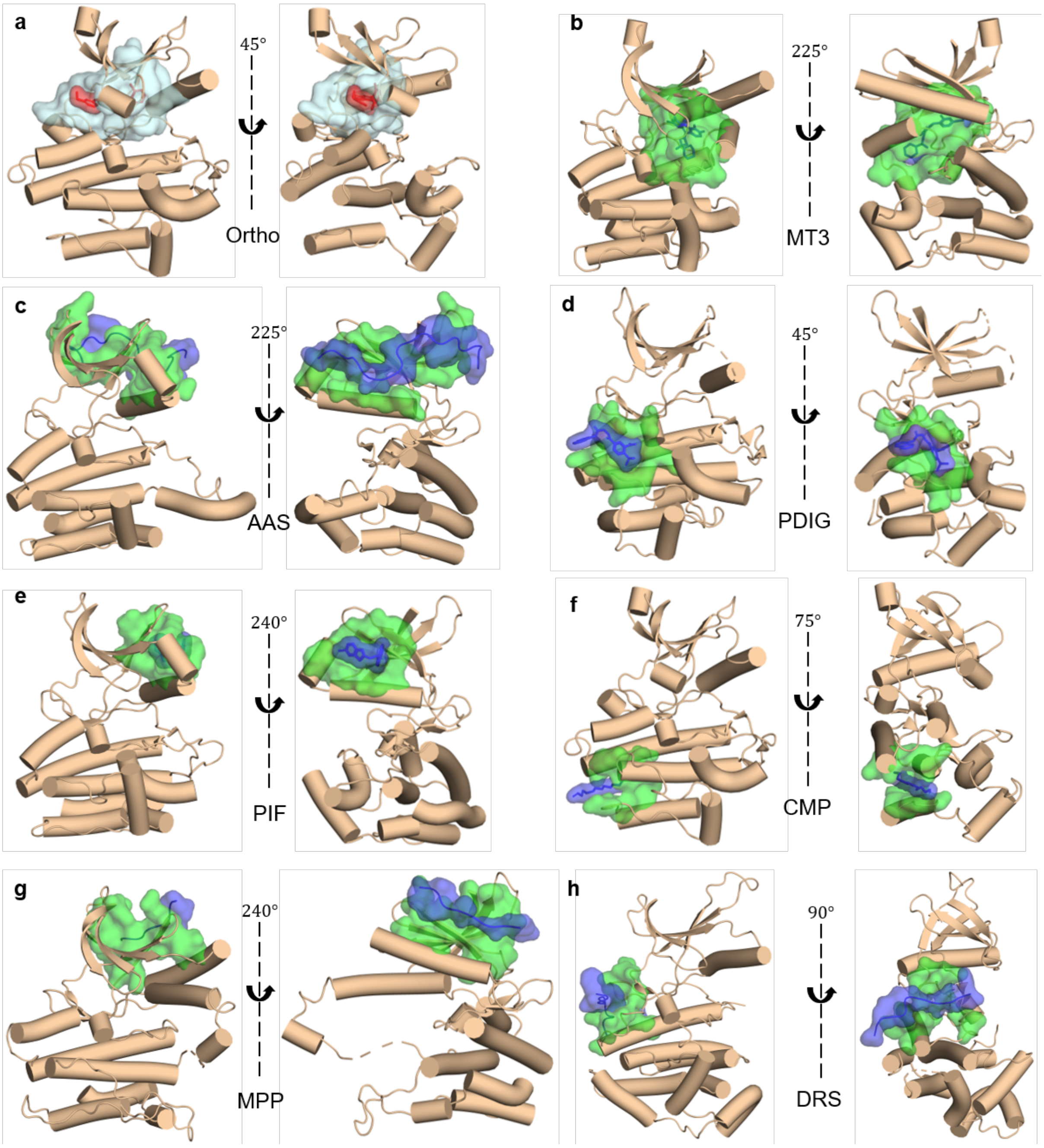


**Figure S8** Three-dimensional representation of orthosteric and allosteric pockets. The orthosteric site (cyan surface) is defined by residues within 5 Å of P16 ligand (red) in ABL1 kinase (PDB id: 1OPL). The green surfaces represent residues within 5 Å of ligand/peptide represented in blue. Two orientations are shown for better visualization. (b) MAP2K1 (PDB id: 4AN2) for MEK1/2 type III inhibitor (MT3) site, (c) AURKA (PDB id: 4C3P) for aurora A activation segment (AAS) site, (d) CHEK1 (PDB id: 3JVS) for PDIG motif site, (e) PDPK1 (PDB id: 4RQK) for PDK1 interacting fragment (PIF) site, (f) ABL1 (PDB id: 1OPL) for c-Abl myristoyl pocket (CMP) site, (g) MAP2K4 (PDB id: 3ALO) for MKK4 p38a peptide (MPP) site and (h) MAPK8 (PDB id: 1UKI) for D-recruitment (DRS) site. The detailed list of allosteric residues for each site is provided in Table S5.

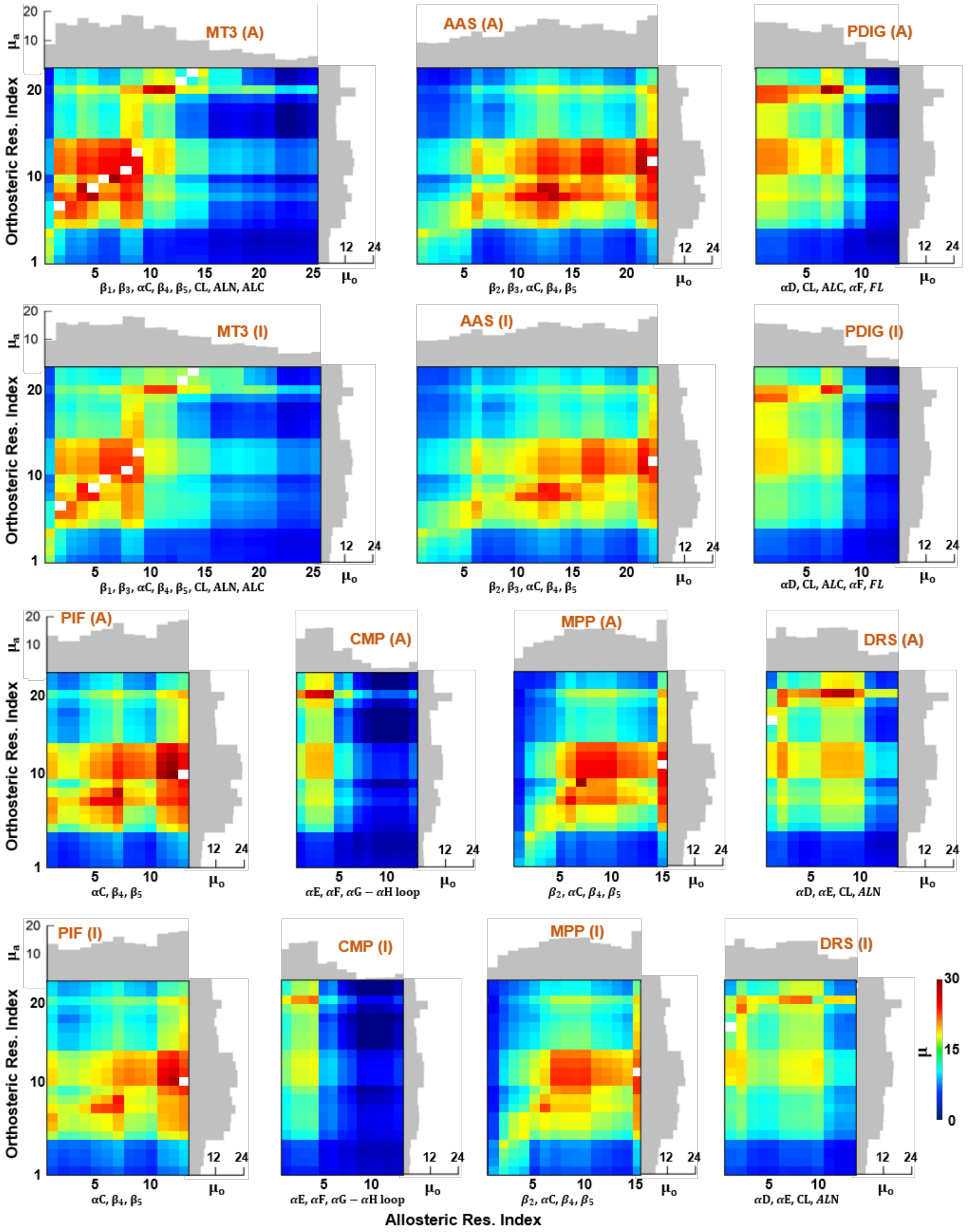


**Figure S9** Effective thermodynamic coupling maps, i.e., $\mu = < \Delta G_c >$, across 104 active-inactive kinase pairs between the orthosteric and allosteric sites. A structure based multiple sequence alignment[2] is employed to identify structurally identical positions corresponding to orthosteric and allosteric sites in all kinases. The "A" within bracket corresponds to active (rows 1 and 3) conformation, while "I" corresponds to inactive kinase conformation (rows 2 and 4). The inactive kinases are always less coupled compared to the active kinases.

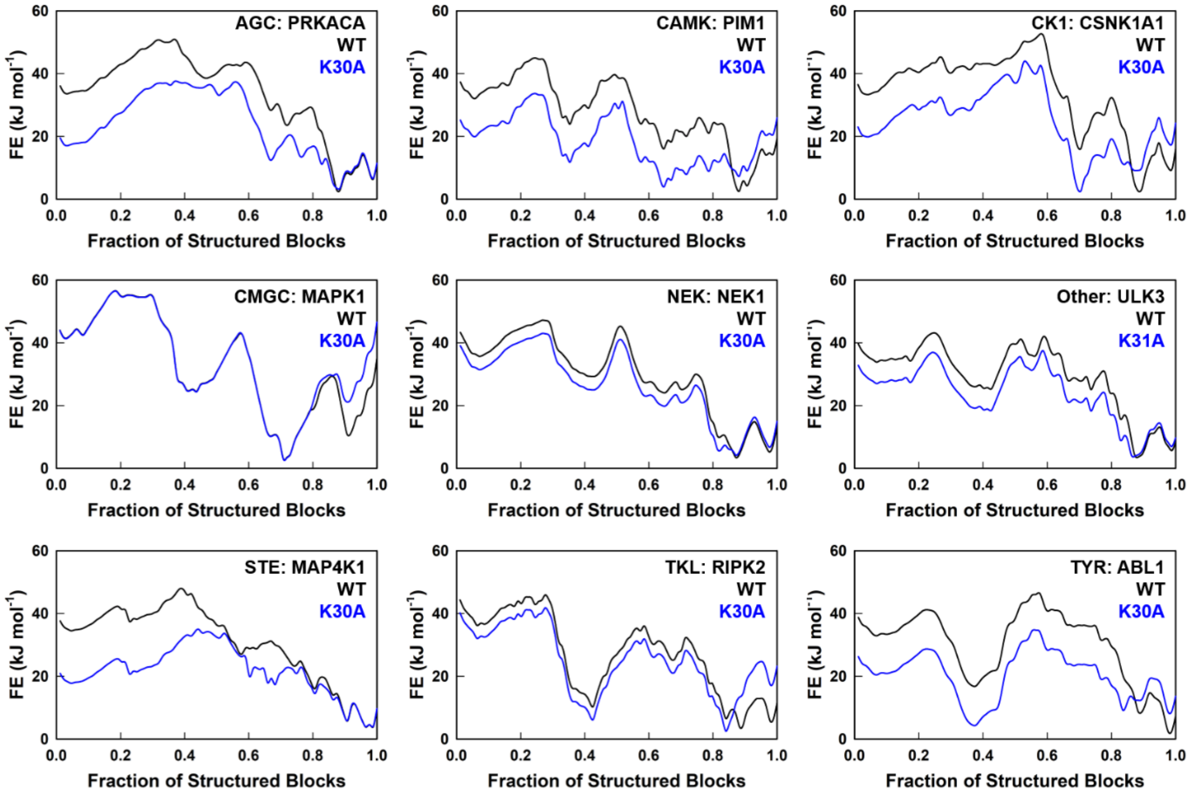


**Figure S10** Free energy profiles of one representative kinase from the nine families before (WT, black) and after mutating a structurally identical position corresponding to K30 in PRKACA from AGC family to alanine (blue). The PDB ids are 3OVV for PRKACA, 1XWS for PIM1, 6GZD for CSNK1A1, 3SA0 for MAPK1, 4APC for NEK1, 6FDY for ULK3, 7M0M for MAP4K1, 6S1F for RIPK2 and 6XR6 for ABL1. RGC family is not included, as there are no experimental structures are available.

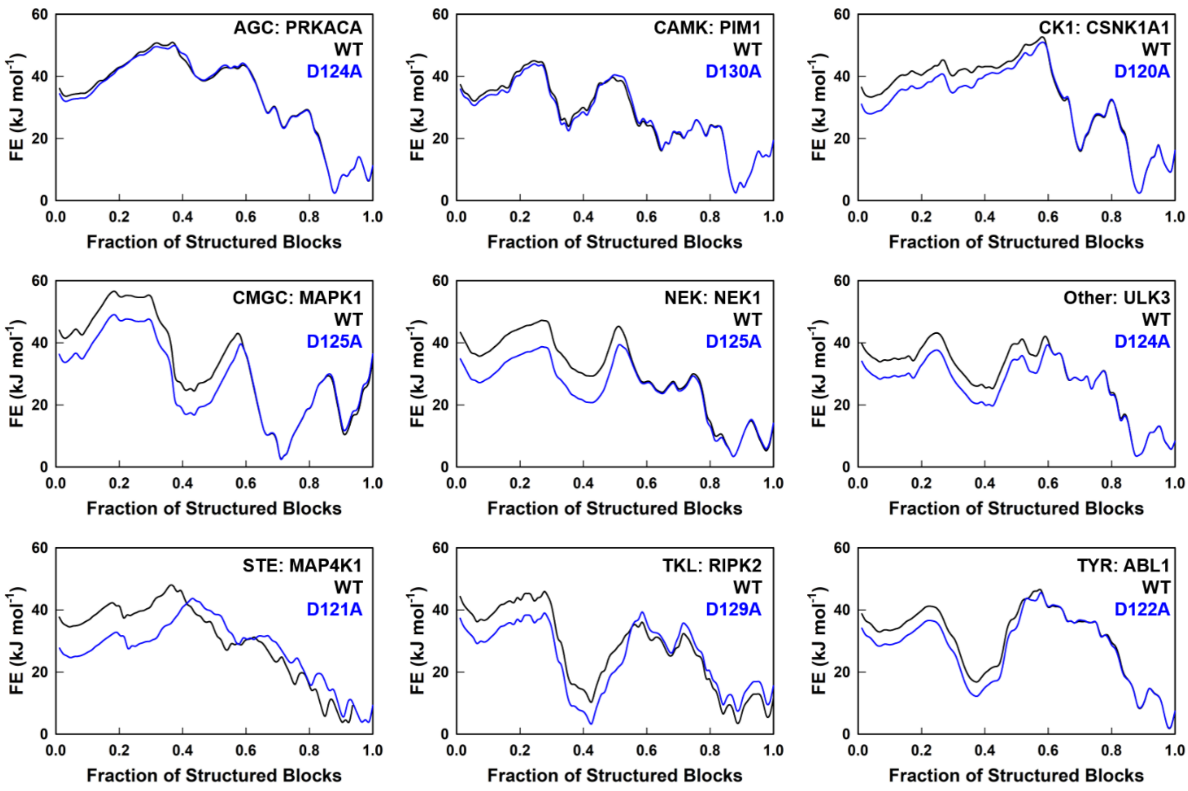


**Figure S11** Same as Figure S10 but on mutating structurally identical position corresponding to D124 (from HRD motif) in PRKACA to alanine.

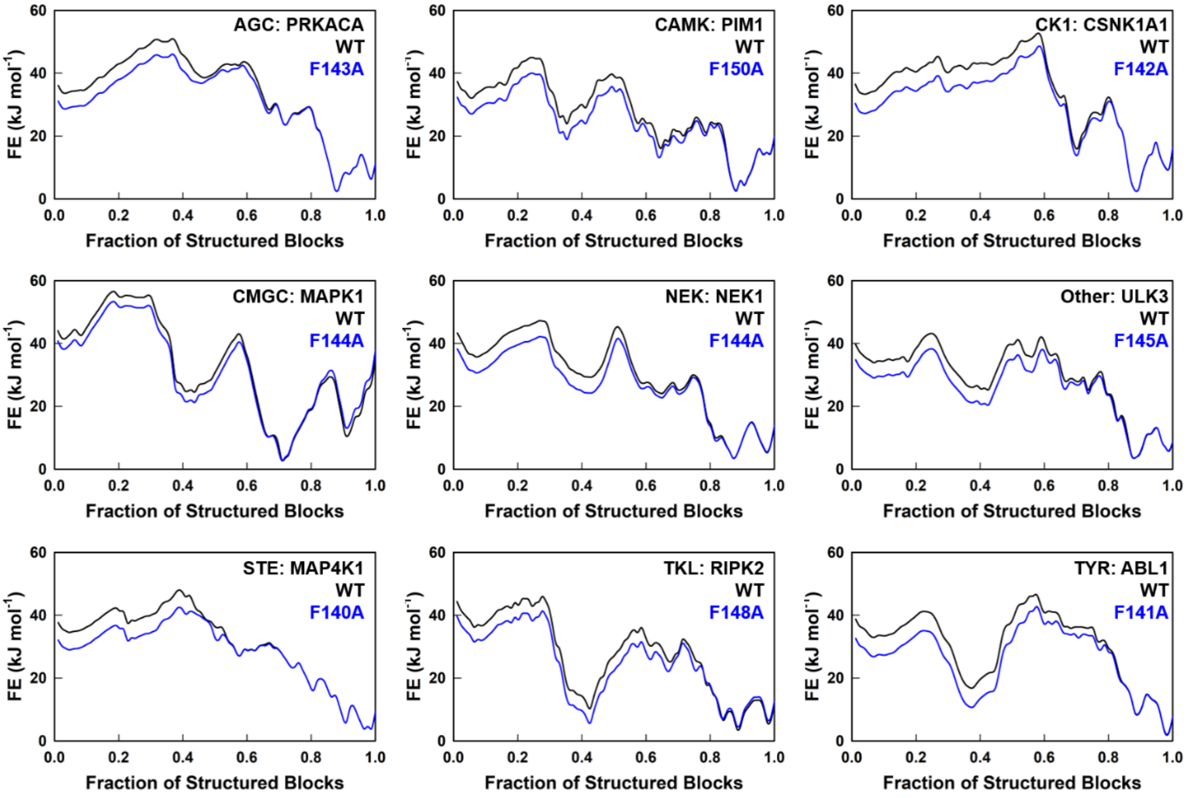


**Figure S12** Same as a Figure S10 but on mutating structurally identical position corresponding to F143 (from DFG motif) in PRKACA to alanine.

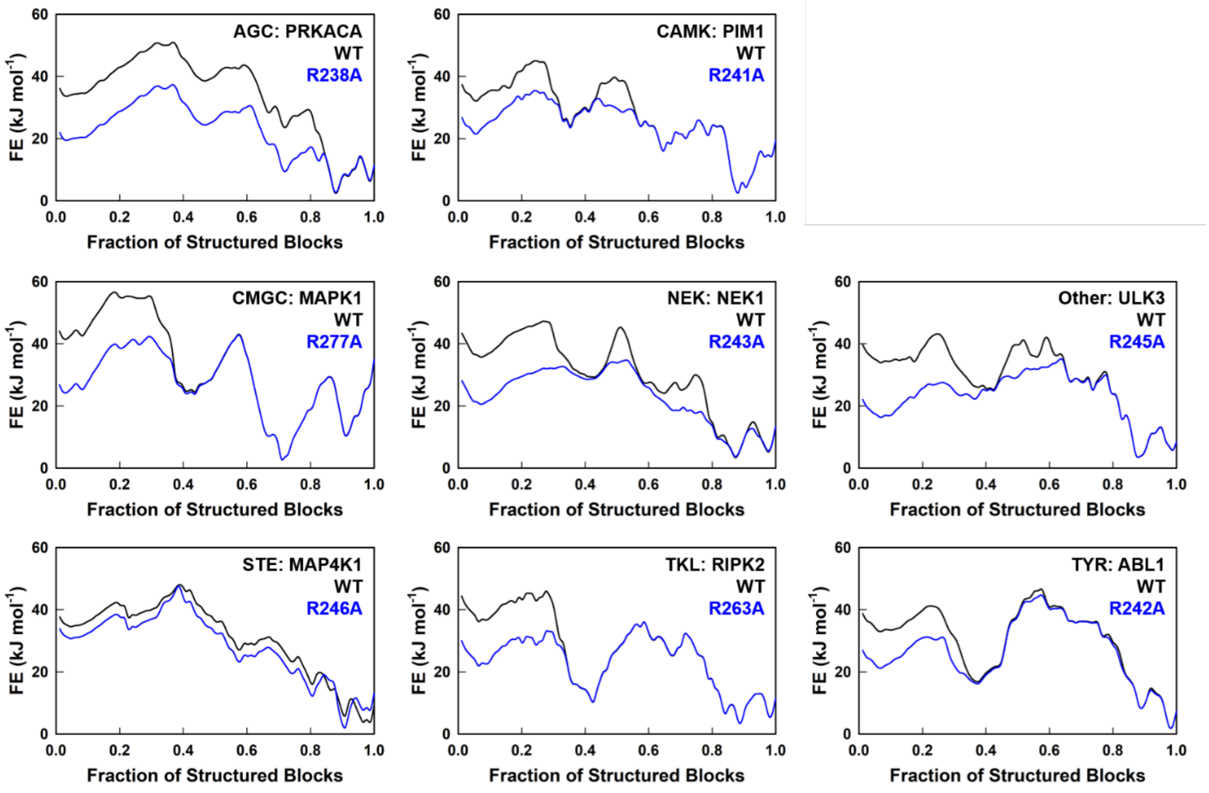


**Figure S13** Same as a Figure S10 but on mutating structurally identical position corresponding to R238 in PRKACA (from $C_t$-lobe) to alanine. CSNK1A1 from CK1 family has an alanine at this structurally identical position (i.e., position 255) instead of arginine, and hence no plot is shown for this kinase.

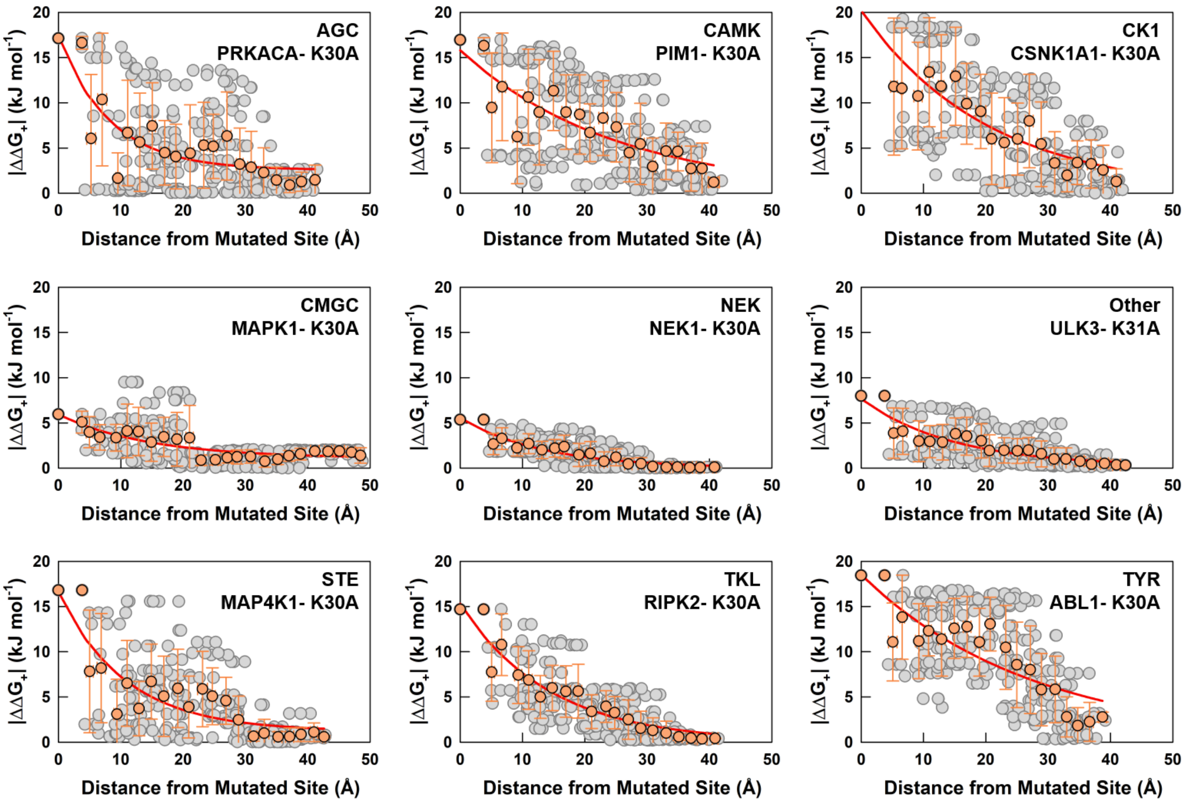


**Figure S14** Absolute mutational response ($|\Delta\Delta G_{+}|$) as a function of distance from the mutated site from one representative kinase from all nine families upon mutating a structurally identical position at $N_t$-lobe corresponding to K30 in PRKACA to alanine. The binned $|\Delta\Delta G_{+}|$ (within 2 Å) is represented by orange points with bars indicating standard deviations of both $|\Delta\Delta G_{+}|$ as well as distance from the mutated site (standard deviations along the distance axis are too small to be seen). The exponential fit to the circles is shown in red from which the coupling distance is extracted.

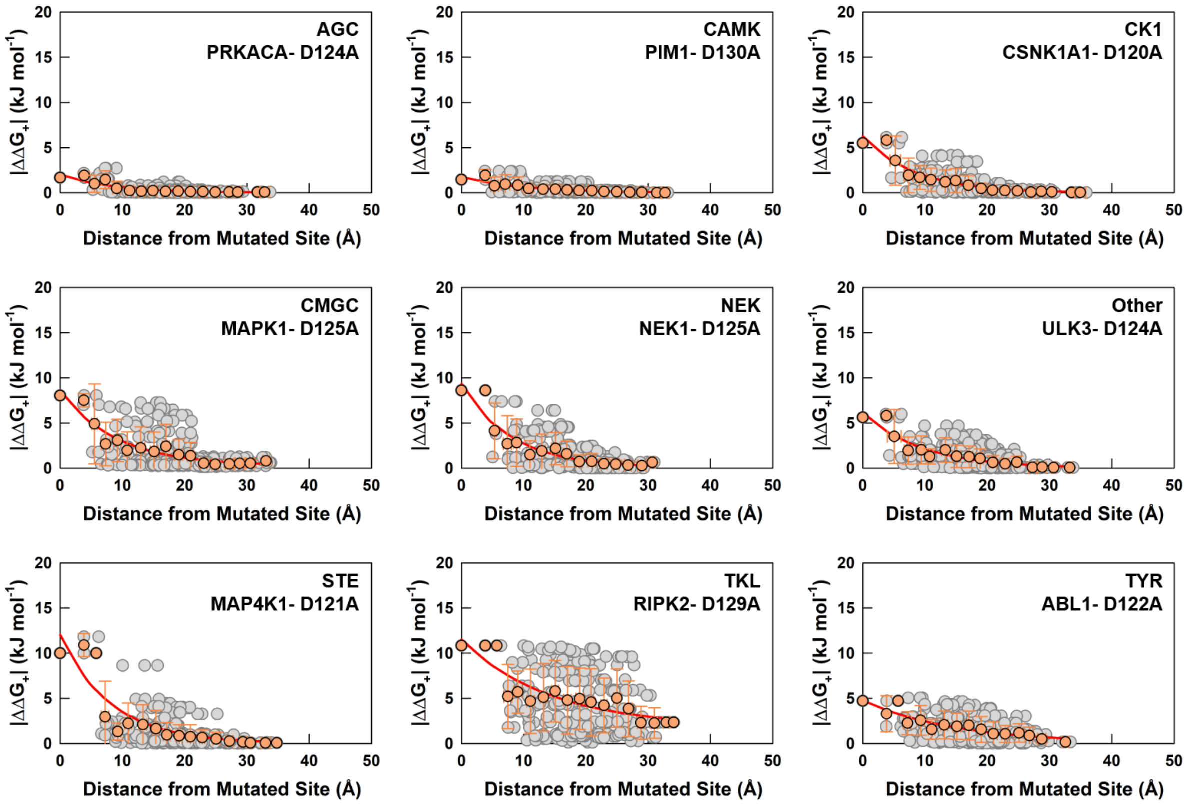


**Figure S15** Same as Figure S14 but on mutating structurally identical positions corresponding to D124 (of HRD motif) in PRKACA to alanine.

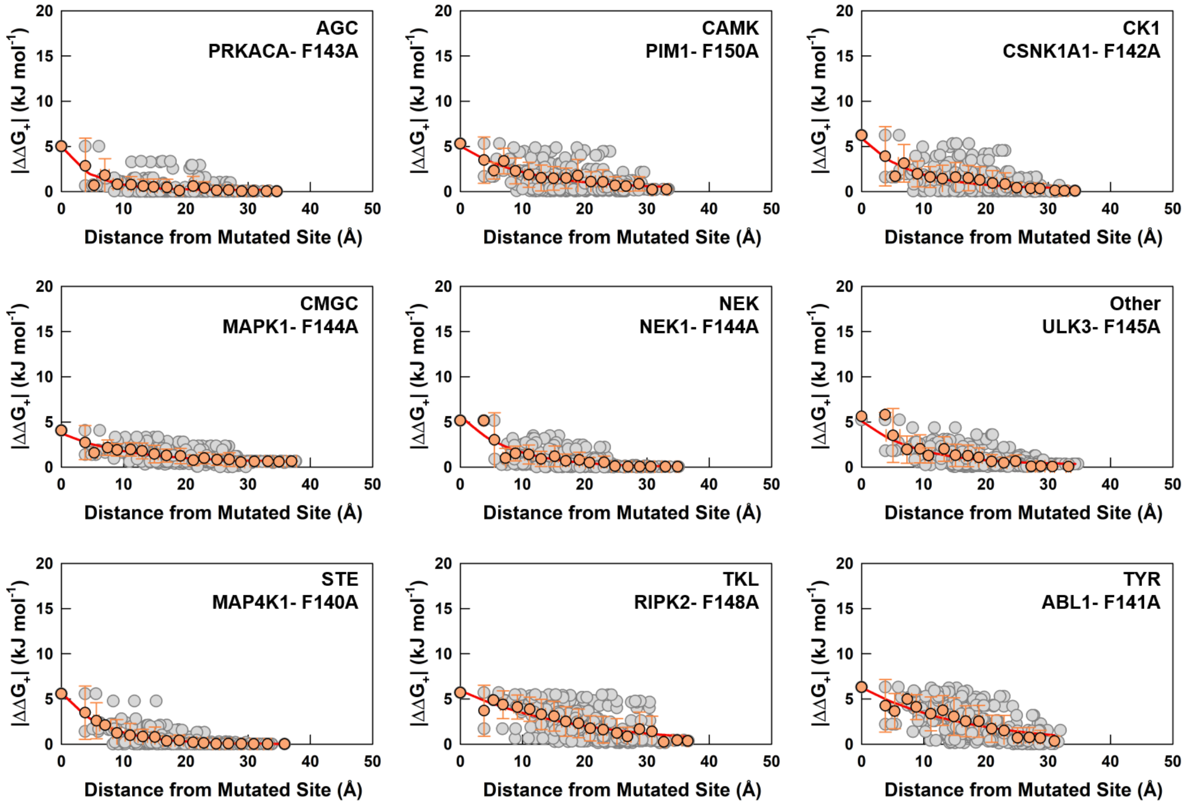


**Figure S16** Same as Figure S14 but on mutating structurally identical positions corresponding to F143 (of DFG motif) in PRKACA to alanine.

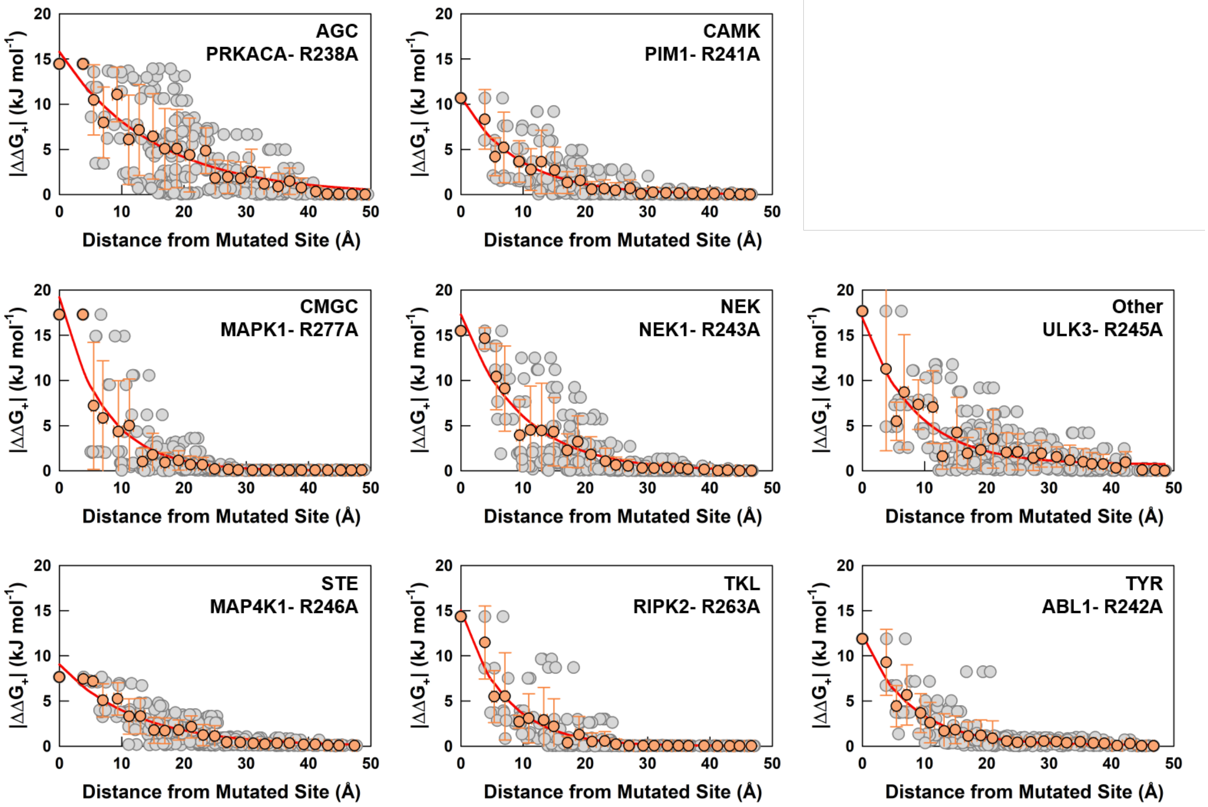


**Figure S17** Same as Figure S14 but on mutating structurally identical positions corresponding to R238 in PRKACA to alanine. CSNK1A1 from CK1 family has an alanine at this structurally identical position (i.e., position 255) instead of arginine, and hence no plot is shown for this kinase.

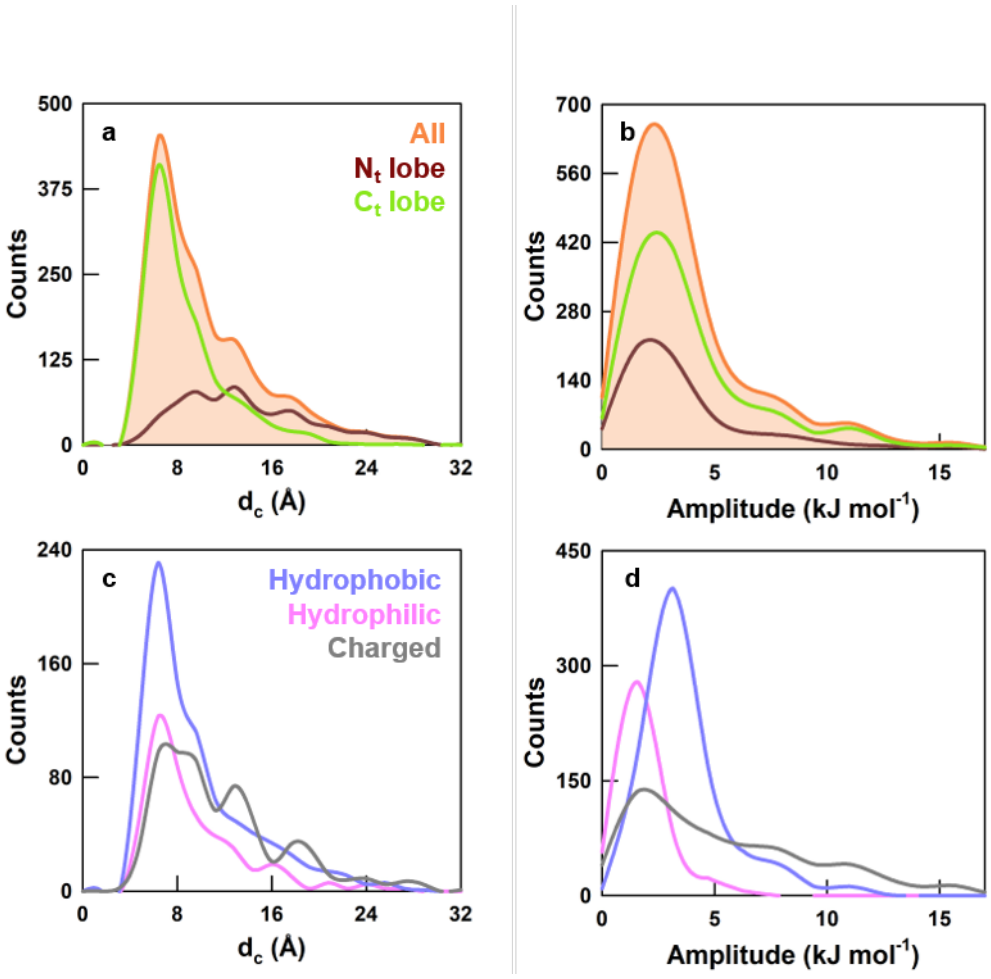


**Figure S18** (a, b) Distribution of coupling distances ($d_c$, panel a) and perturbation amplitudes (amplitude, panel b) for all the mutated residues (orange, 1975 mutations), residues of $N_t$-lobe (brown) and residues of $C_t$-lobe (green). (c, d) The distribution of $d_c$ (panel c) and amplitude (panel d) for all the mutated residues differentiated as hydrophobic (blue), hydrophilic (pink) and charged (gray).

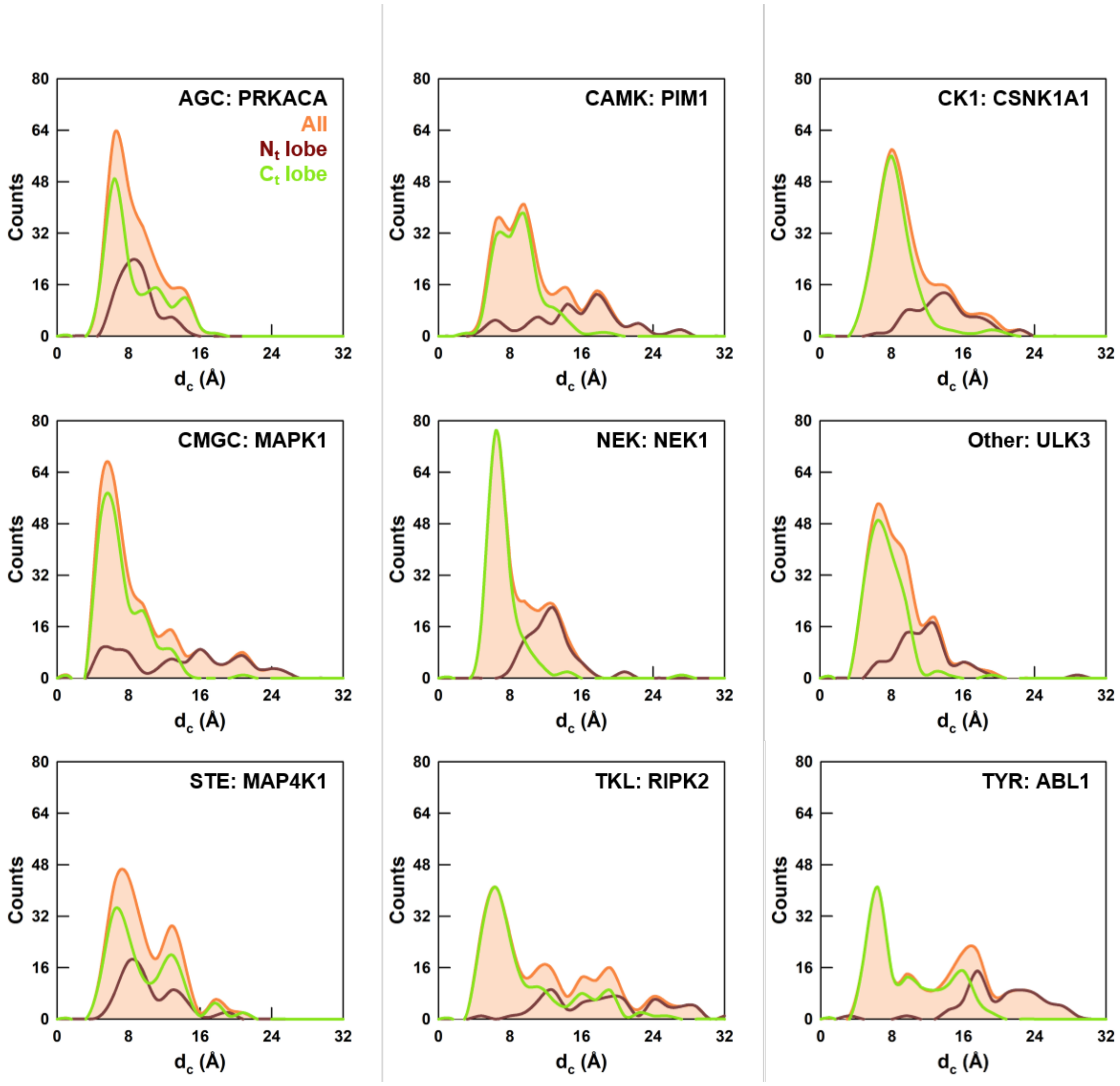


**Figure S19** Distribution of coupling distances ($d_c$) from alanine-scanning mutagenesis for all residues (orange), residues in the $N_t$-lobe (brown) and $C_t$-lobe (green) lobe for one member from each family, except RGC.

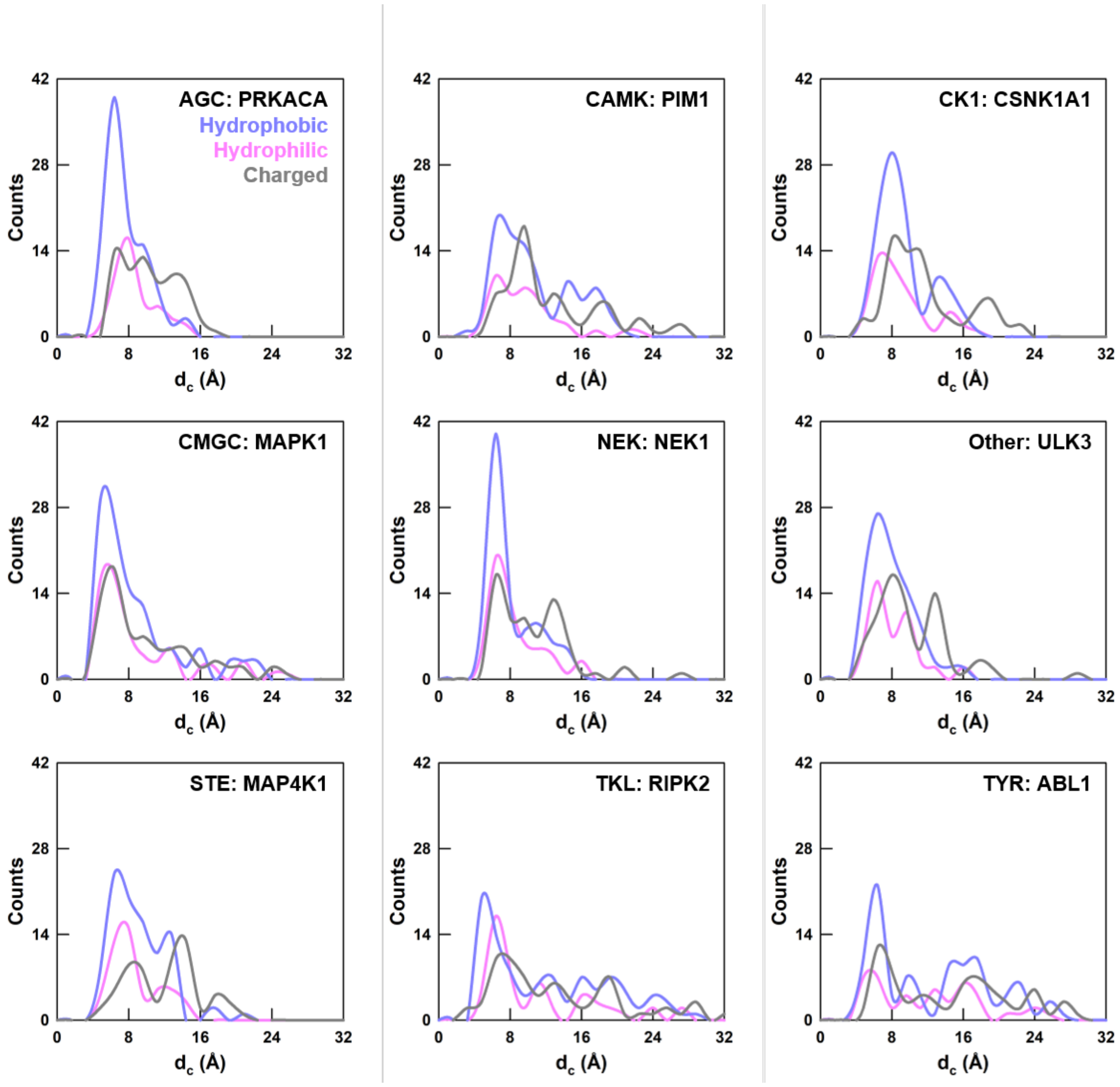


**Figure S20** Distribution of coupling distances ($d_c$) from alanine-scanning mutagenesis for hydrophobic (pink), hydrophilic (blue) and charged (grey) residues.

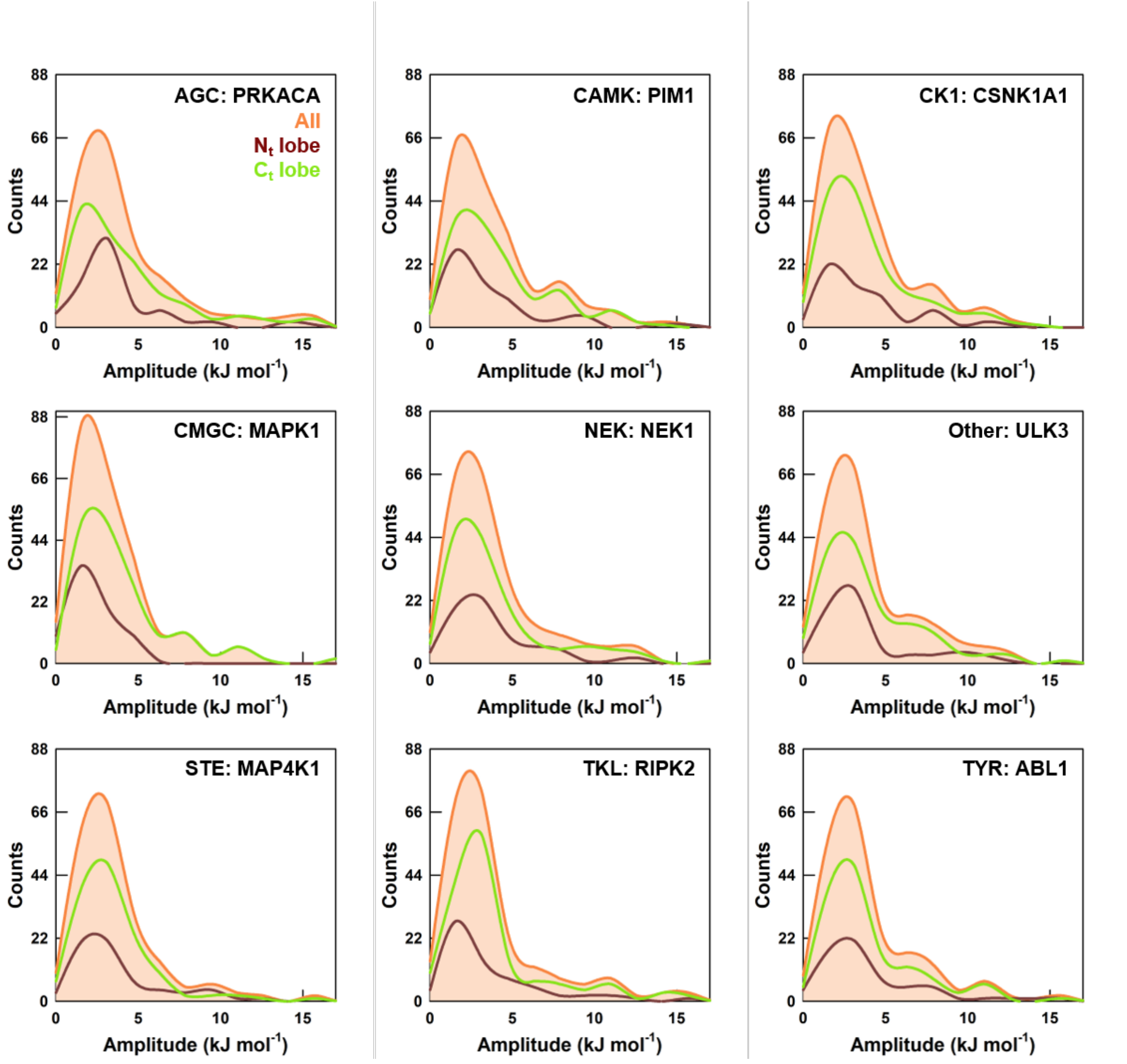


**Figure S21** Distribution of perturbation amplitudes from alanine-scanning mutagenesis of the residues belonging to $N_t$-lobe (brown), and $C_t$-lobe (green). The sum of the two distributions is shown in red.

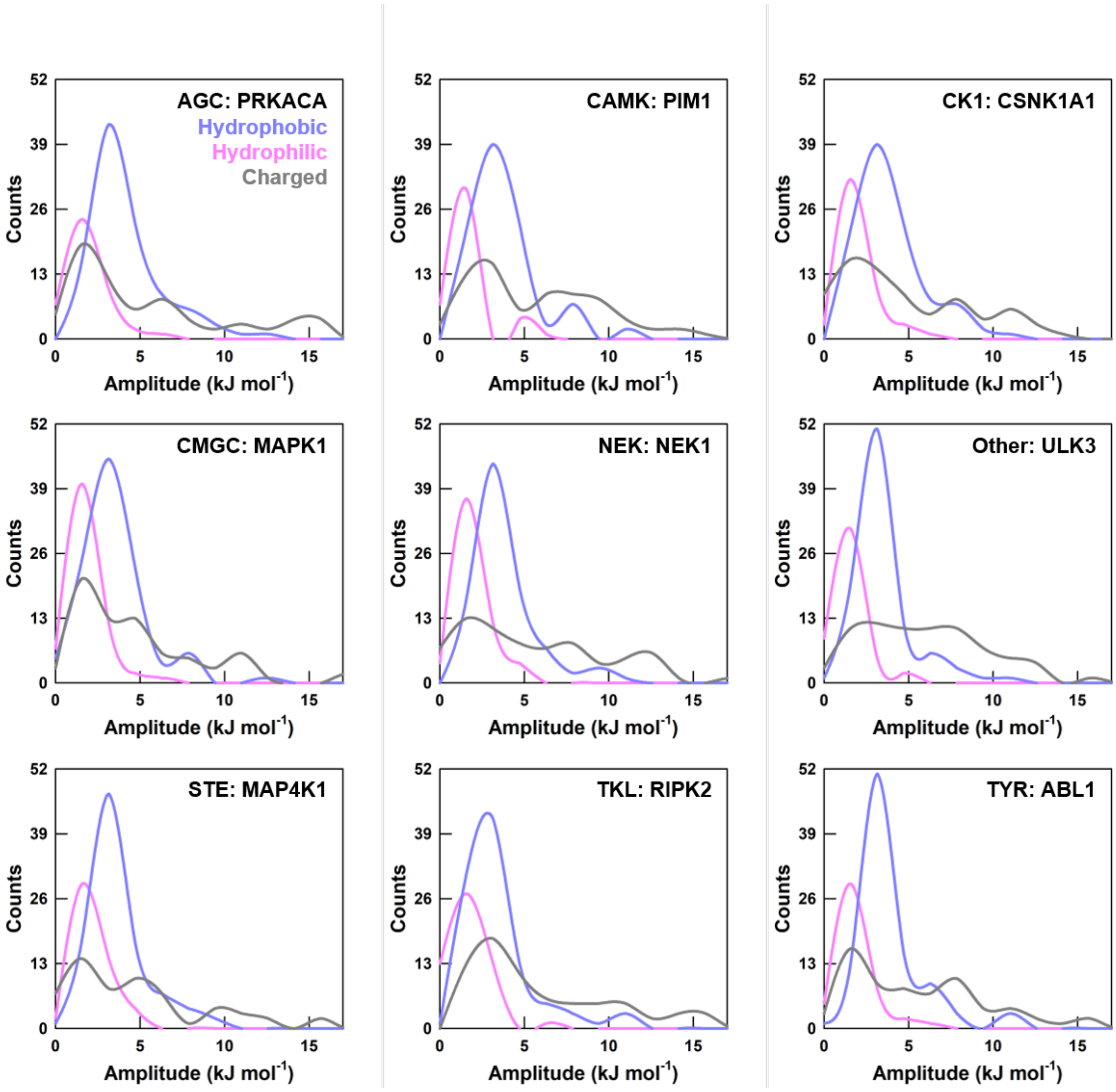


**Figure S22** Distribution of perturbation amplitudes from alanine-scanning mutagenesis of hydrophobic (pink), hydrophilic (blue) and charged (grey) residues.

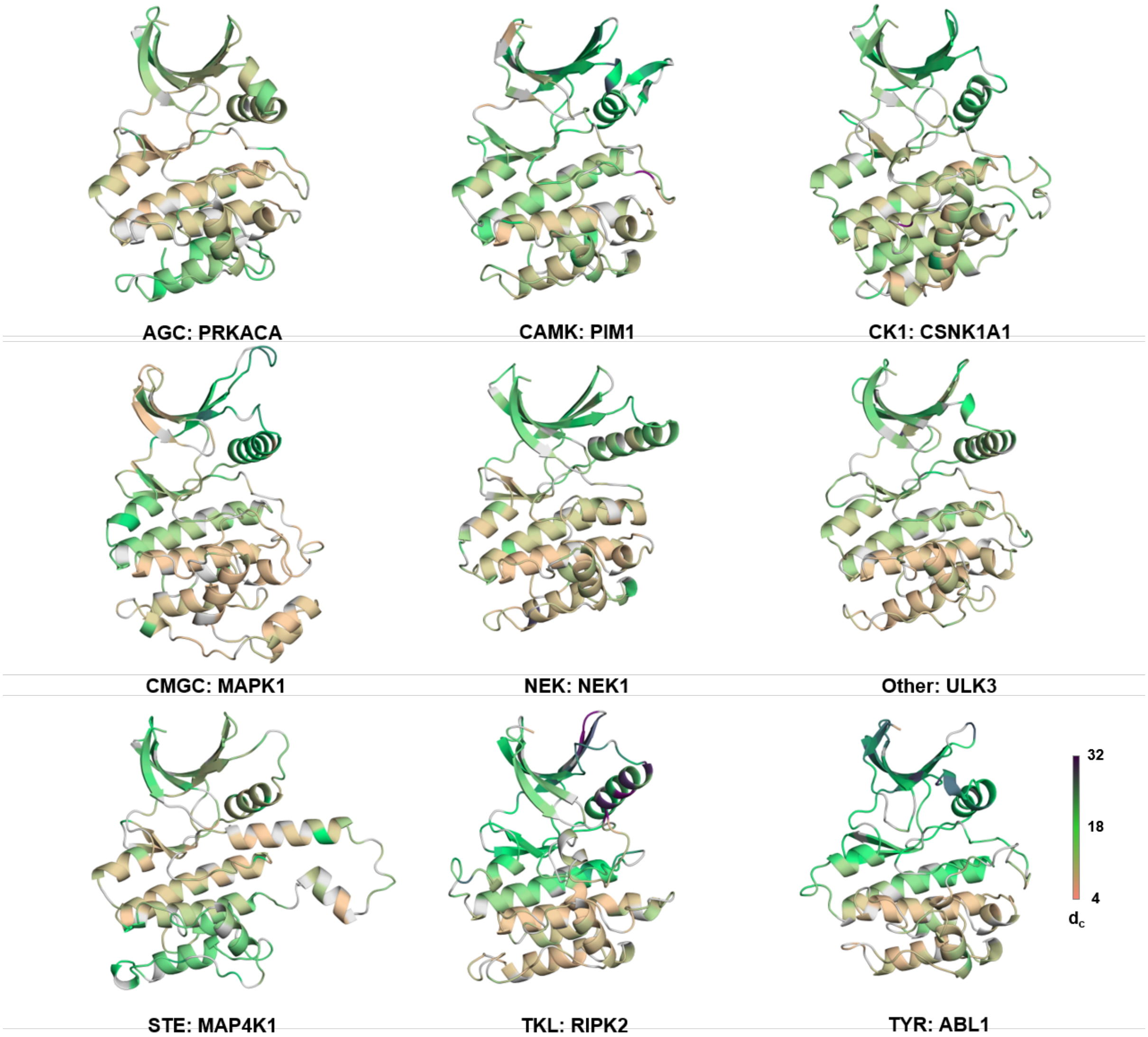


**Figure S23** The magnitude of coupling distance (colorbar at the bottom right in Å) mapped onto one representative structure from each family. Alanine, glycine, and proline residues are colored grey and were not mutated.

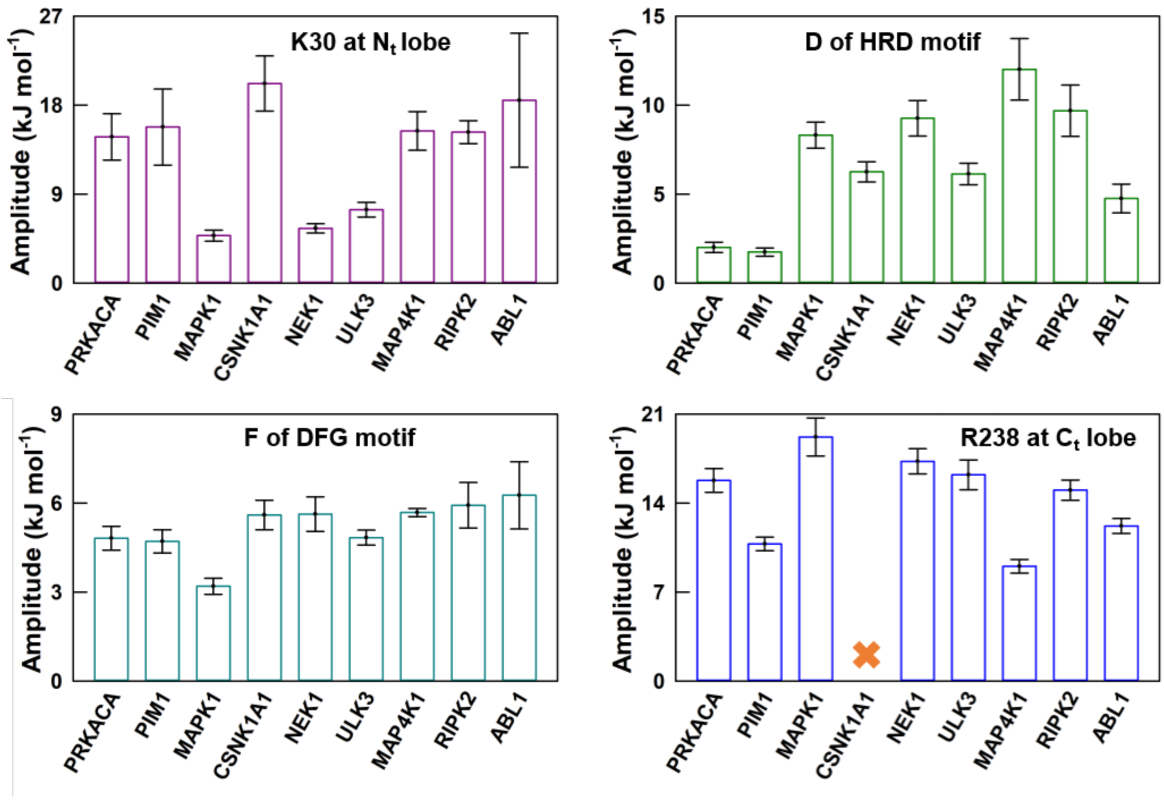


**Figure S24** Perturbation amplitudes corresponding to Figure 6g in the main text. Note that the amplitudes are quite different, highlighting changes in interaction networks not obvious in structural superimpositions.

**Table S1** Statistics of kinases from 10 typical and 6 atypical families as provided int the pkinfam[3] (https://ftp.uniprot.org/pub/databases/uniprot/current_release/knowledgebase/complete/docs/pkinfam.txt). 14 out of the 484 typical kinase genes translate to two catalytic domains and these are the kinase genes of RPS6KA1, RPS6KA2, RPS6KA3, RPS6KA4, RPS6KA5, RPS6KA6 from AGC; OBSCN, SPEG from CAMK; JAK1, JAK2, JAK3, TYK2 from TYR and EIF2AK4, RPS6KC1 from 'Other' family.

| **Kinase Family** | **Number of Kinases** | | **Experimental Structures** | |
|---|---|---|---|---|
| | **Genes** | **Domains** | **Available** | **Used** |
| Typical Protein Kinase | | | | |
| AGC | 58 | 64 | 31 | 30 |
| CAMK | 75 | 77 | 39 | 38 |
| CK1 | 12 | 12 | 11 | 11 |
| CMGC | 61 | 61 | 40 | 38 |
| NEK | 11 | 11 | 3 | 3 |
| RGC | 5 | 5 | 0 | 0 |
| STE | 56 | 56 | 33 | 32 |
| TKL | 34 | 34 | 22 | 22 |
| TYR or TK | 91 | 95 | 68 | 61 |
| Other | 81 | 83 | 43 | 39 |
| Total | 484 | 498 | 290 | 274 |
| Atypical protein kinase | | | | |
| ADCK | 5 | 5 | 1 | 0 |
| Alpha-type | 6 | 6 | 0 | 0 |
| FAST | 1 | 1 | 0 | 0 |
| PDK/BCKDK | 5 | 5 | 4 | 0 |
| PI3/PI4-kinase | 10 | 10 | 8 | 0 |
| RIO-type | 3 | 3 | 2 | 0 |
| Total | 30 | 30 | 15 | 0 |

**Table S2** Excluded kinases.

| S. No. | Kinase Name | PDB id | Deleted residues or Mean TM-score | Exclusion Reason |
|---|---|---|---|---|
| 1 | MASTL | 5LOH | 195-739 | Residues were deleted during structure determination |
| 2 | SRPK1 | 1WAK | 257-473 | |
| 3 | SRPK2 | 5MYV | 257-507 | |
| 4 | EIF2AK4 (domain 2) | 6N3O | 658-780 | |
| 5 | CSF1R | 2OGV | 682-740 | |
| 6 | KDR | 6GQQ | 940-990 | |
| 7 | EIF2AK3 | 4G34 | 661-868 | |
| 8 | PEAK1 | 6BHC | 1409-1451 | |
| 9 | FLT3 | 6IL3 | 711-761 | |
| 10 | KIT | 6MOB | 694-753 | |
| 11 | CDC7 | 6YA7 | 228-345, 467-533 | |
| 12 | PDGFRA | 5K5X | 695-768 | |
| 13 | PBK | 5J0A | 0.62 | Mean TM-score is below 0.7 |
| 14 | TRIB1 | 5CEM | 0.57 | |
| 15 | FLT1 | 3HNG | 0.63 | |
| 16 | ERBB3 | 6OP9 | 0.68 | |

**Table S3** Database of Kinase containing 274 members along with complementary 104 inactive pair for 104 active state kinases. The kinase name with suffix "_2" represents second catalytic domain of the kinase. The "A" within bracket corresponds to active conformation while "I" corresponds to inactive conformation PDB ids. No. of residues, blocks and microstates in the table are for active state kinase.

| Family | Name | PDB ID | UniProt ID | vdW Interaction Energy (J $mol^{-1}$) | No. of Residues | No. of Blocks | No. of Microstates |
|---|---|---|---|---|---|---|---|
| AGC | PDPK1 | 1H1W (A), 3NAX (I) | O15530 | -72.7 | 261 | 88 | 4820591 |
| AGC | AKT2 | 1O6L (A), 1GZN (I) | P31751 | -75 | 258 | 87 | 4626594 |
| AGC | RPS6KA5 | 1VZO (I) | O75582 | -76.5 | 270 | 88 | 4837441 |
| AGC | GRK6 | 2ACX (A) | P43250 | -75.1 | 263 | 90 | 5278639 |
| AGC | PRKCB | 2I0E (A) | P05771 | -79.8 | 259 | 86 | 4390056 |
| AGC | SGK1 | 2R5T (I) | O00141 | -77.7 | 258 | 86 | 4394646 |
| AGC | DMPK | 2VD5 (A) | Q09013 | -76 | 269 | 92 | 5760704 |
| AGC | RPS6KA1 | 2Z7R (I) | Q15418 | -76.5 | 260 | 87 | 4607914 |
| AGC | PRKCI | 3A8X (A), 6ILZ (I) | P41743 | -77.8 | 269 | 93 | 6017373 |
| AGC | PRKCA | 3IW4 (A) | P17252 | -78.6 | 259 | 84 | 3993466 |
| AGC | RPS6KA5_2 | 3KN6 (A) | O75582 | -76.6 | 262 | 88 | 4820624 |
| AGC | PRKACA | 3OVV (A), 4AE6 (I) | P17612 | -73.4 | 255 | 87 | 4607010 |
| AGC | PRKCH | 3TXO (A) | P24723 | -82 | 260 | 87 | 4590362 |
| AGC | ROCK1 | 3V8S (A), 5WNF (I) | Q13464 | -76 | 263 | 88 | 4823182 |
| AGC | RPS6KB1 | 3WF8 (A), 4L3J (I) | P23443 | -79.9 | 262 | 88 | 4817467 |
| AGC | PKN2 | 4CRS (A) | Q16513 | -78.1 | 260 | 86 | 4401662 |
| AGC | AKT1 | 4GV1 (A), 6HHJ (I) | P31749 | -74.2 | 259 | 86 | 4418451 |
| AGC | RPS6KA3_2 | 4JG6 (A), 4D9T (I) | P51812 | -76.6 | 258 | 88 | 4790666 |
| AGC | GRK2 | 4MK0 (A), 3CIK (I) | P25098 | -78.4 | 263 | 89 | 5039210 |
| AGC | RPS6KA1_2 | 4NIF (A), 2WNT (I) | Q15418 | -74.9 | 258 | 88 | 4797511 |
| AGC | RPS6KA3 | 4NUS (I) | P51812 | -76.3 | 260 | 87 | 4605937 |
| AGC | PKN1 | 4OTD (I) | Q16512 | -77.9 | 260 | 87 | 4592975 |
| AGC | GRK5 | 4TND (A), 6PJX (I) | P34947 | -75.9 | 263 | 89 | 5058857 |
| AGC | GRK4 | 4YHJ (A) | P32298 | -74.8 | 263 | 89 | 5060734 |
| AGC | PRKCQ | 5F9E (A) | Q04759 | -75.3 | 255 | 83 | 3809105 |
| AGC | CDC42BPB | 5OTF (A) | Q9Y5S2 | -77.2 | 267 | 91 | 5527351 |
| AGC | PRKG1 | 6BG2 (A) | Q13976 | -76 | 260 | 88 | 4834248 |
| AGC | STK38 | 6BXI (I) | Q15208 | -71 | 294 | 96 | 6862860 |
| AGC | RPS6KA6 | 6G77 (I) | Q9UK32 | -79.4 | 258 | 88 | 4830772 |

| | | | | | | | |
|---|---|---|---|---|---|---|---|
| AGC | ROCK2 | 7JNT (A), 4L6Q (I) | O75116 | -75.3 | 263 | 89 | 5046417 |
| CAMK | CHEK1 | 1IA8 (A) | O14757 | -75.5 | 257 | 88 | 4825478 |
| CAMK | TTN | 1TKI (I) | Q8WZ42 | -78.1 | 255 | 88 | 4817269 |
| CAMK | PIM1 | 1XWS (A), 6KZI (I) | P11309 | -73.7 | 253 | 85 | 4171640 |
| CAMK | DAPK2 | 2A2A (A) | Q9UIK4 | -77.4 | 263 | 91 | 5504308 |
| CAMK | MKNK2 | 2AC3 (I) | Q9HBH9 | -77.2 | 305 | 101 | 8346457 |
| CAMK | CHEK2 | 2CN5 (A), 3I6U (I) | O96017 | -78.7 | 267 | 91 | 5504637 |
| CAMK | CAMK1G | 2JAM (I) | Q96NX5 | -81.2 | 255 | 88 | 4789339 |
| CAMK | CAMK1D | 2JC6 (I) | Q8IU85 | -85.2 | 257 | 89 | 5034696 |
| CAMK | MAPKAPK2 | 2OZA (A) | P49137 | -76.9 | 262 | 90 | 5210267 |
| CAMK | MARK3 | 2QNJ (I) | P27448 | -78.3 | 252 | 85 | 4202053 |
| CAMK | CAMK2G | 2V7O (A) | Q13555 | -77.1 | 259 | 92 | 5748173 |
| CAMK | CAMK2D | 2VN9 (A) | Q13557 | -77.1 | 259 | 90 | 5258517 |
| CAMK | CAMK2A | 2VZ6 (A), 3SOA (I) | Q9UQM7 | -77 | 259 | 90 | 5259318 |
| CAMK | DAPK1 | 2W4J (A), 1WVY (I) | P53355 | -75.7 | 263 | 89 | 5020464 |
| CAMK | CAMK4 | 2W4O (I) | Q16566 | -77 | 255 | 88 | 4829885 |
| CAMK | STK11 | 2WTK (A) | Q15831 | -78.4 | 261 | 89 | 5041219 |
| CAMK | MYLK4 | 2X4F (I) | Q86YV6 | -76.9 | 256 | 87 | 4611897 |
| CAMK | PHKG2 | 2Y7J (A) | P15735 | -74.1 | 268 | 92 | 5751221 |
| CAMK | CAMK2B | 3BHH (A) | Q13554 | -75.8 | 259 | 90 | 5265133 |
| CAMK | DAPK3 | 3BHY (A) | O43293 | -76.6 | 263 | 89 | 5042655 |
| CAMK | CASK | 3C0G (A) | O14936 | -73.1 | 265 | 92 | 5734775 |
| CAMK | PASK | 3DLS (A) | Q96RG2 | -78.8 | 253 | 88 | 4765986 |
| CAMK | MARK2 | 3IEC (A), 5EAK (I) | Q7KZI7 | -77.4 | 252 | 85 | 4198972 |
| CAMK | MAPKAPK3 | 3R1N (A) | Q16644 | -76.2 | 261 | 89 | 4991836 |
| CAMK | CAMK1 | 4FG7 (A), 4FGB (I) | Q14012 | -81 | 257 | 87 | 4587604 |
| CAMK | PIM2 | 4X7Q (A) | Q9P1W9 | -78.7 | 255 | 85 | 4180667 |
| CAMK | PRKAA2 | 4ZHX (A), 2H6D (I) | P54646 | -78.7 | 253 | 85 | 4173504 |
| CAMK | MARK4 | 5ES1 (I) | Q96L34 | -76.3 | 252 | 83 | 3818885 |
| CAMK | DCLK1 | 5JZJ (A), 7F3G (I) | O15075 | -78.6 | 258 | 87 | 4599787 |
| CAMK | STK40 | 5L2Q (A) | Q8N2I9 | -81.6 | 297 | 102 | 8710038 |
| CAMK | MELK | 5TWU (A), 5K00 (I) | Q14680 | -78.9 | 253 | 85 | 4176144 |
| CAMK | MKNK1 | 5WVD (I) | Q9BUB5 | -76.2 | 326 | 108 | 10932349 |
| CAMK | SNRK | 5YKS (I) | Q9NRH2 | -74.7 | 254 | 82 | 3622296 |
| CAMK | MARK1 | 6C9D (A), 2HAK (I) | Q9P0L2 | -77 | 252 | 87 | 4610502 |
| CAMK | PRKAA1 | 6C9H (A), 6C9F (I) | Q13131 | -76.4 | 253 | 86 | 4375860 |
| CAMK | STK17B | 6Y6H (A) | O94768 | -77.5 | 261 | 91 | 5495269 |

| | | | | | | | |
|---|---|---|---|---|---|---|---|
| CAMK | STK17A | 7QUE (A), 7QUF (I) | Q9UEE5 | -78.3 | 261 | 91 | 5519981 |
| CAMK | TRIB2 | 7UPM (I) | Q92519 | -77.2 | 248 | 82 | 3614420 |
| CK1 | CSNK1G2 | 2C47 (A) | P78368 | -72.6 | 270 | 93 | 6033606 |
| CK1 | CSNK1G3 | 2CHL (A) | Q9Y6M4 | -72 | 271 | 95 | 6572626 |
| CK1 | CSNK1G1 | 2CMW (A) | Q9HCP0 | -74 | 272 | 93 | 6028089 |
| CK1 | VRK3 | 2JII (A) | Q8IV63 | -75.3 | 292 | 99 | 7682146 |
| CK1 | VRK2 | 2V62 (A), 5UU1 (I) | Q86Y07 | -74 | 291 | 101 | 8356275 |
| CK1 | CSNK1E | 4HOK (A), 4HNI (I) | P49674 | -87.5 | 269 | 92 | 5752061 |
| CK1 | TTBK1 | 4NFN (A), 7Q8W (I) | Q5TCY1 | -79 | 264 | 91 | 5529295 |
| CK1 | VRK1 | 6BU6 (A), 2KUL (I) | Q99986 | -72.1 | 281 | 97 | 7138586 |
| CK1 | CSNK1A1 | 6GZD (A) | P48729 | -76.3 | 269 | 95 | 6557437 |
| CK1 | CSNK1D | 6RCH (A) | P48730 | -75.6 | 269 | 94 | 6272903 |
| CK1 | TTBK2 | 6U0K (A), 7Q8Z (I) | Q6IQ55 | -80.1 | 264 | 88 | 4830446 |
| CMGC | MAPK12 | 1CM8 (A), 7CGA (I) | P53778 | -79.5 | 285 | 97 | 7122797 |
| CMGC | CDK2 | 1FIN (A), 4FKU (I) | P24941 | -72.6 | 283 | 97 | 7137526 |
| CMGC | CDK5 | 1H4L (A) | Q00535 | -73.8 | 283 | 97 | 7130597 |
| CMGC | GSK3B | 1J1B (A), 6Y9S (I) | P49841 | -75.5 | 285 | 96 | 6828848 |
| CMGC | MAPK14 | 1WBW (A), 3KQ7 (I) | Q16539 | -78.6 | 285 | 95 | 6491136 |
| CMGC | CLK3 | 2EU9 (A) | P49761 | -73.3 | 317 | 112 | 12744662 |
| CMGC | CDK6 | 2F2C (A), 1BLX (I) | Q00534 | -78.6 | 288 | 95 | 6553517 |
| CMGC | MAPK8 | 2H96 (A), 4QTD (I) | P45983 | -77.4 | 296 | 102 | 8697883 |
| CMGC | DYRK1A | 2VX3 (A), 6UIP (I) | Q13627 | -73.5 | 321 | 111 | 12263790 |
| CMGC | MAPK3 | 2ZOQ (A) | P27361 | -76.9 | 279 | 94 | 6276384 |
| CMGC | MAPK11 | 3GP0 (I) | Q15759 | -78.7 | 285 | 98 | 7401874 |
| CMGC | MAPK9 | 3NPC (I) | P45984 | -76 | 296 | 100 | 8038923 |
| CMGC | MAPK1 | 3SA0 (A), 1WZY (I) | P28482 | -75 | 289 | 96 | 6836936 |
| CMGC | CDKL3 | 3ZDU (A) | Q8IVW4 | -76.2 | 283 | 96 | 6836720 |
| CMGC | CDKL1 | 4AGU (A) | Q00532 | -69 | 284 | 94 | 6285043 |
| CMGC | CDKL2 | 4BBM (I) | Q92772 | -74 | 284 | 96 | 6859869 |
| CMGC | CDKL5 | 4BGQ (A) | O76039 | -73 | 285 | 97 | 7147400 |
| CMGC | CDK12 | 4CXA (A), 7NXK (I) | Q9NYV4 | -81.6 | 294 | 100 | 8078317 |
| CMGC | MAPK10 | 4H39 (A), 4KKE (I) | P53779 | -77 | 296 | 102 | 8709457 |

| | | | | | | | |
|---|---|---|---|---|---|---|---|
| CMGC | MAPK13 | 4MYG (A), 4EYJ (I) | O15264 | -78.1 | 284 | 98 | 7354584 |
| CMGC | CDK9 | 4OR5 (A) | P50750 | -82.3 | 297 | 100 | 8071475 |
| CMGC | CDK1 | 4Y72 (A), 4YC6 (I) | P06493 | -78.1 | 284 | 100 | 8065137 |
| CMGC | MAPK7 | 5BYZ (A) | Q13164 | -76.5 | 293 | 100 | 7993169 |
| CMGC | CDK16 | 5G6V (I) | Q00536 | -75.8 | 282 | 96 | 6852801 |
| CMGC | CDK8 | 5XS2 (A), 5BNJ (I) | P49336 | -74.8 | 315 | 102 | 8759308 |
| CMGC | DYRK3 | 5Y86 (A) | O43781 | -74.9 | 314 | 110 | 11856825 |
| CMGC | CLK2 | 6FYL (A) | P49760 | -71.2 | 317 | 110 | 11861324 |
| CMGC | CLK4 | 6FYV (A) | Q9HAZ1 | -73.3 | 317 | 110 | 11829957 |
| CMGC | DYRK2 | 6HDP (A) | Q92630 | -74.7 | 314 | 108 | 11001495 |
| CMGC | PRPF4B | 6PK6 (A) | Q13523 | -72.9 | 320 | 110 | 11788679 |
| CMGC | MAPK6 | 6YLL (A), 6YLC (I) | Q16659 | -78.3 | 297 | 100 | 8043411 |
| CMGC | CLK1 | 6ZLN (A), 6KHD (I) | P49759 | -71.7 | 317 | 110 | 11873223 |
| CMGC | CDK7 | 7B5Q (A), 1UA2 (I) | P50613 | -76.9 | 284 | 97 | 7109185 |
| CMGC | HIPK2 | 7NCF (A) | Q9H2X6 | -71.7 | 329 | 113 | 13215907 |
| CMGC | CDK13 | 7NXJ (A) | Q14004 | -80 | 294 | 102 | 8752292 |
| CMGC | HIPK3 | 7O7I (A), 7O7J (I) | Q9H422 | -75.5 | 329 | 113 | 13198116 |
| CMGC | CDK4 | 7SJ3 (A), 3G33 (I) | P11802 | -73.5 | 290 | 96 | 6798835 |
| CMGC | CDK11B | 7UKZ (I) | P21127 | -77.3 | 286 | 95 | 6559804 |
| NEK | NEK2 | 2W5A (I) | P51955 | -75.6 | 264 | 88 | 4821001 |
| NEK | NEK7 | 2WQN (I) | Q8TDX7 | -73.2 | 266 | 90 | 5280787 |
| NEK | NEK1 | 4APC (I) | Q96PY6 | -75.6 | 255 | 86 | 4396298 |
| STE | MAP2K2 | 1S9I (I) | P36507 | -78 | 298 | 101 | 8376362 |
| STE | PAK6 | 2C30 (A) | Q9NQU5 | -78.7 | 252 | 86 | 4403935 |
| STE | PAK5 | 2F57 (A) | Q9P286 | -79 | 252 | 88 | 4810479 |
| STE | PAK4 | 2J0I (A), 4FIG (I) | O96013 | -79.4 | 252 | 88 | 4822616 |
| STE | SLK | 2J51 (A), 8BEM (I) | Q9H2G2 | -77.8 | 259 | 89 | 5038071 |
| STE | STK25 | 2XIK (A), 4NZW (I) | O00506 | -76.8 | 251 | 88 | 4837869 |
| STE | STK24 | 3A7H (A), 4O27 (I) | Q9Y6E0 | -75.5 | 251 | 87 | 4619176 |
| STE | MAP2K4 | 3ALO (I) | P45985 | -77 | 266 | 91 | 5516317 |
| STE | OXSR1 | 3DAK (I) | O95747 | -78.7 | 275 | 92 | 5722120 |
| STE | STK26 | 3GGF (I) | Q9P289 | -80.2 | 251 | 85 | 4197499 |
| STE | STRADA | 3GNI (A) | Q7RTN6 | -76.8 | 311 | 107 | 10539886 |
| STE | MAP2K6 | 3VN9 (I) | P52564 | -74.6 | 262 | 88 | 4817573 |
| STE | MAP3K14 | 4IDV (A), 4IDT (I) | Q99558 | -78.6 | 256 | 87 | 4578572 |
| STE | MAP2K1 | 4LMN (I) | Q02750 | -76.5 | 294 | 96 | 6845505 |

| | | | | | | | |
|---|---|---|---|---|---|---|---|
| STE | PAK1 | 4O0R (A), 4ZJI (I) | Q13153 | -78.3 | 252 | 89 | 5006487 |
| STE | MAP4K4 | 4OBP (A), 4U3Y (I) | O95819 | -75.3 | 265 | 89 | 5040662 |
| STE | MAP3K9 | 4UY9 (I) | P80192 | -79 | 269 | 89 | 5041883 |
| STE | MAP3K21 | 4UYA (I) | Q5TCX8 | -77.3 | 278 | 94 | 6243816 |
| STE | MAP3K12 | 5CEN (I) | Q12852 | -76.7 | 242 | 82 | 3573190 |
| STE | STK3 | 5DH3 (A), 4LG4 (I) | Q13188 | -77.4 | 252 | 86 | 4376129 |
| STE | MAP3K7 | 5GJF (I) | O43318 | -82.3 | 256 | 86 | 4361490 |
| STE | MAP3K8 | 5IU2 (A) | P41279 | -78.7 | 251 | 86 | 4394319 |
| STE | MAP4K3 | 5J5T (A) | Q8IVH8 | -80.9 | 258 | 89 | 5024778 |
| STE | MAP3K5 | 5VIL (A), 2CLQ (I) | Q99683 | -73.8 | 259 | 88 | 4819746 |
| STE | TAOK3 | 6BDN (A) | Q9H2K8 | -76.7 | 254 | 86 | 4328594 |
| STE | PAK3 | 6FD3 (A) | O75914 | -77.8 | 252 | 88 | 4794552 |
| STE | STK10 | 6HXF (A), 6EIM (I) | O94804 | -75 | 259 | 89 | 5055665 |
| STE | MAP3K20 | 6JUU (I) | Q9NYL2 | -73.9 | 262 | 90 | 5252693 |
| STE | TNIK | 6RA7 (A), 5D7A (I) | Q9UKE5 | -76.5 | 265 | 92 | 5766970 |
| STE | STK4 | 6YAT (A) | Q13043 | -76.5 | 252 | 85 | 4179846 |
| STE | MAP2K7 | 6YG1 (A), 6QFL (I) | O14733 | -73.3 | 261 | 89 | 5032653 |
| STE | MAP4K1 | 7M0M (A), 7M0K (I) | Q92918 | -82.4 | 258 | 88 | 4791597 |
| TKL | ACVR2B | 2QLU (A) | Q13705 | -74.2 | 291 | 98 | 7453568 |
| TKL | BMPR2 | 3G2F (A) | Q13873 | -76.5 | 302 | 104 | 9489242 |
| TKL | BMPR1B | 3MDY (A) | O00238 | -74.2 | 291 | 99 | 7754102 |
| TKL | ACVRL1 | 3MY0 (A) | P37023 | -79.4 | 291 | 98 | 7426270 |
| TKL | RAF1 | 3OMV (A) | P04049 | -80.8 | 261 | 88 | 4803922 |
| TKL | ILK | 3REP (I) | Q13418 | -72.8 | 254 | 88 | 4765983 |
| TKL | LIMK1 | 3S95 (A), 7ATU (I) | P53667 | -75 | 266 | 90 | 5275971 |
| TKL | ACVR2A | 3SOC (A) | P27037 | -75.2 | 294 | 98 | 7447335 |
| TKL | ACVR1 | 4C02 (A), 6T6D (I) | Q04771 | -79.2 | 295 | 100 | 8056416 |
| TKL | BRAF | 4MNE (A), 5VAM (I) | P15056 | -78.1 | 261 | 87 | 4564861 |
| TKL | LIMK2 | 4TPT (I) | P53671 | -75.5 | 278 | 93 | 6030701 |
| TKL | TNNI3K | 4YFI (A), 7MGJ (I) | Q59H18 | -78.3 | 261 | 92 | 5723211 |
| TKL | TGFBR1 | 5E8S (A) | P36897 | -75.9 | 291 | 97 | 7146465 |
| TKL | TGFBR2 | 5E8V (A), 5E8Y (I) | P37173 | -74.9 | 301 | 101 | 8419725 |
| TKL | IRAK1 | 6BFN (A) | P51617 | -71.7 | 310 | 106 | 10098736 |
| TKL | RIPK1 | 6C4D (I) | Q13546 | -76.8 | 273 | 92 | 5785917 |
| TKL | IRAK4 | 6LXY (A), 6EGF (I) | Q9NWZ3 | -80.6 | 269 | 93 | 6003630 |
| TKL | IRAK3 | 6RUU (A) | Q9Y616 | -76.2 | 288 | 99 | 7724646 |

| | | | | | | | |
|---|---|---|---|---|---|---|---|
| TKL | RIPK2 | 6S1F (A), 5NG3 (I) | O43353 | -74.1 | 277 | 94 | 6271578 |
| TKL | KSR2 | 7JUR (I) | Q6VAB6 | -77.7 | 266 | 90 | 5264564 |
| TKL | KSR1 | 7JUW (I) | Q8IVT5 | -75.1 | 271 | 89 | 5043464 |
| TKL | LRRK2 | 7LI4 (I) | Q5S007 | -80.4 | 260 | 88 | 4802868 |
| TYR | TEK | 1FVR (I) | Q02763 | -76.7 | 273 | 89 | 5030512 |
| TYR | TNK2 | 1U46 (A), 4EWH (I) | Q07912 | -70.4 | 260 | 89 | 5027247 |
| TYR | ZAP70 | 1U59 (A), 2OZO (I) | P43403 | -73.3 | 263 | 92 | 5767716 |
| TYR | SRC | 1Y57 (A), 2SRC (I) | P12931 | -75.2 | 254 | 87 | 4595152 |
| TYR | FYN | 2DQ7 (A) | P06241 | -75 | 254 | 89 | 5023935 |
| TYR | PTK2 | 2ETM (A), 4K9Y (I) | Q05397 | -73.2 | 259 | 88 | 4802961 |
| TYR | HCK | 2HK5 (A), 5ZJ6 (I) | P08631 | -73.1 | 254 | 88 | 4823161 |
| TYR | FGFR2 | 2PSQ (A), 8E1X (I) | P21802 | -76.6 | 290 | 97 | 7131825 |
| TYR | EPHA3 | 2QOC (A), 3DZQ (I) | P29320 | -76.4 | 262 | 89 | 5058645 |
| TYR | EPHA5 | 2R2P (A) | P54756 | -76.6 | 262 | 89 | 5055999 |
| TYR | EPHA7 | 2REI (A), 3DKO (I) | Q15375 | -77.7 | 262 | 90 | 5286146 |
| TYR | ABL2 | 2XYN (A), 3HMI (I) | P42684 | -76.6 | 252 | 84 | 3989821 |
| TYR | LYN | 3A4O (I) | P07948 | -75 | 255 | 88 | 4813586 |
| TYR | ERBB4 | 3BCE (A), 2R4B (I) | Q15303 | -74 | 268 | 94 | 6241947 |
| TYR | FES | 3BKB (A) | P07332 | -83.6 | 262 | 90 | 5278358 |
| TYR | INSR | 3BU3 (A), 1IRK (I) | P06213 | -80 | 276 | 96 | 6837931 |
| TYR | PTK2B | 3CC6 (I) | Q14289 | -72 | 259 | 89 | 5051057 |
| TYR | CSK | 3D7U (A), 1BYG (I) | P41240 | -80.6 | 255 | 87 | 4609142 |
| TYR | FGFR1 | 3GQI (A), 3TT0 (I) | P11362 | -79.2 | 290 | 100 | 8046354 |
| TYR | EPHA8 | 3KUL (A) | P29322 | -82.2 | 262 | 89 | 5042387 |
| TYR | LCK | 3LCK (A), 6PDJ (I) | P06239 | -79.4 | 254 | 87 | 4601800 |
| TYR | TYK2_2 | 3NYX (A), 6VNS (I) | P29597 | -76 | 280 | 96 | 6839177 |
| TYR | MST1R | 3PLS (I) | Q04912 | -77 | 264 | 91 | 5488824 |
| TYR | ERBB2 | 3PP0 (I) | P04626 | -73.9 | 268 | 92 | 5746548 |
| TYR | ITK | 3QGY (A), 4M14 (I) | Q08881 | -77.4 | 253 | 87 | 4569026 |
| TYR | IGF1R | 3QQU (A), 3NW5 (I) | P08069 | -79.5 | 276 | 94 | 6266773 |
| TYR | BMX | 3SXS (I) | P51813 | -76.5 | 259 | 85 | 4187738 |

| | | | | | | | |
|---|---|---|---|---|---|---|---|
| TYR | EPHB4 | 3ZEW (A), 6FNI (I) | P54760 | -79.4 | 285 | 98 | 7418882 |
| TYR | EPHB2 | 3ZFM (I) | P29323 | -76.1 | 264 | 91 | 5510986 |
| TYR | MET | 3ZZE (I) | P08581 | -80.6 | 268 | 90 | 5277310 |
| TYR | ROR2 | 3ZZW (I) | Q01974 | -77.8 | 274 | 89 | 5018777 |
| TYR | NTRK2 | 4AT5 (I) | Q16620 | -74.9 | 270 | 91 | 5508356 |
| TYR | SYK | 4GFG (A), 4FL2 (I) | P43405 | -74.8 | 261 | 88 | 4812599 |
| TYR | JAK1 | 4L00 (I) | P23458 | -72.7 | 273 | 91 | 5533462 |
| TYR | FGFR4 | 4TYE (I) | P22455 | -81.7 | 289 | 100 | 8061115 |
| TYR | ROS1 | 4UXL (I) | P08922 | -74 | 278 | 94 | 6265923 |
| TYR | EGFR | 4WKQ (A), 5U8L (I) | P00533 | -72.9 | 268 | 91 | 5485227 |
| TYR | NTRK3 | 4YMJ (I) | Q16288 | -75.2 | 302 | 101 | 8381245 |
| TYR | JAK1_2 | 5KHW (A), 6TPF (I) | P23458 | -71.6 | 279 | 95 | 6559888 |
| TYR | EPHB3 | 5L6P (I) | P54753 | -76.7 | 264 | 92 | 5768664 |
| TYR | JAK3_2 | 5LWM (A), 7C3N (I) | P52333 | -71.9 | 290 | 102 | 8731003 |
| TYR | EPHB1 | 5MJB (A) | P54762 | -76.5 | 264 | 89 | 5056919 |
| TYR | BTK | 5P9I (I) | Q06187 | -72.7 | 254 | 86 | 4390329 |
| TYR | JAK2_2 | 5TQ8 (A), 4IVA (I) | O60674 | -70.3 | 276 | 95 | 6573763 |
| TYR | AXL | 5U6B (A) | P30530 | -76.2 | 272 | 92 | 5775270 |
| TYR | JAK2 | 5UT3 (I) | O60674 | -73.3 | 265 | 90 | 5280730 |
| TYR | PTK6 | 6CZ2 (I) | Q13882 | -74.8 | 255 | 87 | 4596978 |
| TYR | NTRK1 | 6D22 (I) | P04629 | -74.1 | 272 | 91 | 5500827 |
| TYR | DDR2 | 6FER (I) | Q16832 | -74 | 287 | 95 | 6562689 |
| TYR | DDR1 | 6FIO (I) | Q08345 | -77.1 | 296 | 101 | 8374332 |
| TYR | LMTK3 | 6SEQ (I) | Q96Q04 | -71.7 | 279 | 93 | 6005907 |
| TYR | ROR1 | 6TU9 (I) | Q01973 | -78 | 274 | 95 | 6544438 |
| TYR | RYK | 6TUA (I) | P34925 | -76 | 274 | 93 | 6011612 |
| TYR | PTK7 | 6VG3 (I) | Q13308 | -75.8 | 271 | 92 | 5777509 |
| TYR | ABL1 | 6XR6 (A), 6XRG (I) | P00519 | -75.4 | 252 | 85 | 4190330 |
| TYR | MERTK | 7AAZ (A), 7AW4 (I) | Q12866 | -76 | 272 | 92 | 5775682 |
| TYR | TYK2 | 7AX4 (I) | P29597 | -77.1 | 287 | 94 | 6238508 |
| TYR | ALK | 7BTT (I) | Q9UM73 | -72.6 | 277 | 94 | 6252594 |
| TYR | FGFR3 | 7DHL (A), 6PNX (I) | P22607 | -81.4 | 290 | 100 | 8042412 |
| TYR | RET | 7JU6 (A) | P07949 | -81 | 293 | 100 | 8061866 |
| TYR | EPHA2 | 8BIN (A), 5NKG (I) | P29317 | -77.6 | 263 | 89 | 5047727 |
| Other | STK16 | 2BUJ (A) | O75716 | -75 | 274 | 93 | 5979730 |
| Other | PLK1 | 2YAC (A) | P53350 | -74.2 | 253 | 84 | 4000624 |
| Other | PLK4 | 3COK (A), 4JXF (I) | O00444 | -76.8 | 254 | 85 | 4179858 |
| Other | CSNK2A1 | 3WAR (A) | P68400 | -71.8 | 286 | 97 | 7156155 |
| Other | AURKB | 4AF3 (I) | Q96GD4 | -71.8 | 251 | 84 | 4003630 |

| | | | | | | | |
|---|---|---|---|---|---|---|---|
| Other | PLK3 | 4B6L (A) | Q9H4B4 | -68.7 | 253 | 85 | 4184181 |
| Other | IKBKB | 4E3C (A) | O14920 | -84.9 | 286 | 95 | 6526316 |
| Other | TBK1 | 4EUU (A), 4IWO (I) | Q9UHD2 | -76.6 | 302 | 102 | 8714572 |
| Other | STK32A | 4FR4 (A) | Q8WU08 | -77.9 | 259 | 86 | 4391064 |
| Other | PLK2 | 4I5P (A) | Q9NYY3 | -74.2 | 253 | 86 | 4396194 |
| Other | RNASEL | 4OAU (A) | Q05823 | -78 | 222 | 76 | 2678610 |
| Other | HASPIN | 4QTC (A) | Q8TF76 | -77.1 | 315 | 105 | 9782212 |
| Other | BMP2K | 4W9W (A), 5I3R (I) | Q9NSY1 | -73.4 | 266 | 91 | 5505085 |
| Other | AAK1 | 4WSQ (A) | Q2M2I8 | -74.5 | 270 | 94 | 6269295 |
| Other | GAK | 4Y8D (A), 4C57 (I) | O14976 | -76.6 | 275 | 92 | 5755543 |
| Other | BUB1 | 5DMZ (A) | O43683 | -75.6 | 299 | 103 | 9039570 |
| Other | CHUK | 5EBZ (A) | O15111 | -82.5 | 288 | 98 | 7423439 |
| Other | TLK2 | 5O0Y (A) | Q86UE8 | -74.1 | 280 | 98 | 7433747 |
| Other | WNK3 | 5O21 (A), 5O2C (I) | Q9BYP7 | -73.7 | 259 | 89 | 5068536 |
| Other | AURKA | 5OS5 (A), 2J4Z (I) | O14965 | -73.8 | 251 | 84 | 4006857 |
| Other | WNK1 | 5TF9 (A) | Q9H4A3 | -76.9 | 259 | 86 | 4409181 |
| Other | WEE1 | 5V5Y (A) | P30291 | -78.2 | 271 | 94 | 6292654 |
| Other | PKMYT1 | 5VCW (A) | Q99640 | -78.2 | 250 | 84 | 3988332 |
| Other | WEE2 | 5VDK (A) | P0C1S8 | -79.3 | 275 | 92 | 5750115 |
| Other | PRAG1 | 5VE6 (I) | Q86YV5 | -74.9 | 352 | 114 | 13387046 |
| Other | CAMKK1 | 6CD6 (A) | Q8N5S9 | -77.6 | 282 | 98 | 7436997 |
| Other | EIF2AK2 | 6D3K (A), 2A1A (I) | P19525 | -75.7 | 272 | 94 | 6313561 |
| Other | ULK3 | 6FDY (A), 6FDZ (I) | Q6PHR2 | -74.5 | 257 | 87 | 4593563 |
| Other | AURKC | 6GR8 (I) | Q9UQB9 | -72 | 251 | 83 | 3819747 |
| Other | TTK | 6GVJ (I) | P33981 | -76.4 | 267 | 92 | 5764869 |
| Other | ULK1 | 6QAS (A) | O75385 | -77.6 | 263 | 86 | 4364522 |
| Other | ULK2 | 6QAV (A) | Q8IYT8 | -81.2 | 263 | 85 | 4154043 |
| Other | ULK4 | 6TSZ (I) | Q96C45 | -74.5 | 277 | 94 | 6296761 |
| Other | ERN1 | 6W3C (A), 4Z7H (I) | O75460 | -80.3 | 262 | 86 | 4380409 |
| Other | TP53RK | 6WQX (A) | Q96S44 | -74.6 | 221 | 73 | 2273286 |
| Other | CSNK2A2 | 7A22 (A), 3E3B (I) | P19784 | -72.5 | 286 | 97 | 7157944 |
| Other | PIK3R4 | 7BL1 (I) | Q99570 | -84.1 | 299 | 100 | 8078492 |
| Other | MLKL | 7JXU (I) | Q8NB16 | -75.1 | 276 | 95 | 6575239 |
| Other | CAMKK2 | 5UY6 (A) | Q96RR4 | -76.7 | 282 | 96 | 6847620 |

**Table S4** Index of the kinases utilized for generating a two-dimensional pseudo-color plot of $\Pi = ln(p_{Ct}/p_{Nt})$. The suffix "_2" represents the second catalytic domain of that kinase.

| Index | AGC | CAMK | CK1 | CMGC | NEK | Other | STE | TKL | TYR |
|---|---|---|---|---|---|---|---|---|---|
| 1 | PDPK1 | CHEK1 | CSNK1G2 | MAPK12 | NEK2 | STK16 | MAP2K2 | ACVR2B | TEK |
| 2 | AKT2 | TTN | CSNK1G3 | CDK2 | NEK7 | PLK1 | PAK6 | BMPR2 | TNK2 |
| 3 | RPS6KA5 | PIM1 | CSNK1G1 | CDK5 | NEK1 | PLK4 | PAK5 | BMPR1B | ZAP70 |
| 4 | GRK6 | DAPK2 | VRK3 | GSK3B | | CSNK2A1 | PAK4 | ACVRL1 | SRC |
| 5 | PRKCB | MKNK2 | VRK2 | MAPK14 | | AURKB | SLK | RAF1 | FYN |
| 6 | SGK1 | CHEK2 | CSNK1E | CLK3 | | PLK3 | STK25 | ILK | PTK2 |
| 7 | DMPK | CAMK1G | TTBK1 | CDK6 | | IKBKB | STK24 | LIMK1 | HCK |
| 8 | RPS6KA1 | CAMK1D | VRK1 | MAPK8 | | TBK1 | MAP2K4 | ACVR2A | FGFR2 |
| 9 | PRKCI | MAPKAPK2 | CSNK1A1 | DYRK1A | | STK32A | OXSR1 | ACVR1 | EPHA3 |
| 10 | PRKCA | MARK3 | CSNK1D | MAPK3 | | PLK2 | STK26 | BRAF | EPHA5 |
| 11 | RPS6KA5_2 | CAMK2G | TTBK2 | MAPK11 | | RNASEL | STRADA | LIMK2 | EPHA7 |
| 12 | PRKACA | CAMK2D | | MAPK9 | | HASPIN | MAP2K6 | TNNI3K | ABL2 |
| 13 | PRKCH | CAMK2A | | MAPK1 | | BMP2K | MAP3K14 | TGFBR1 | LYN |
| 14 | ROCK1 | DAPK1 | | CDKL3 | | AAK1 | MAP2K1 | TGFBR2 | ERBB4 |
| 15 | RPS6KB1 | CAMK4 | | CDKL1 | | GAK | PAK1 | IRAK1 | FES |
| 16 | PKN2 | STK11 | | CDKL2 | | BUB1 | MAP4K4 | RIPK1 | INSR |
| 17 | AKT1 | MYLK4 | | CDKL5 | | CHUK | MAP3K9 | IRAK4 | PTK2B |
| 18 | RPS6KA3_2 | PHKG2 | | CDK12 | | TLK2 | MAP3K21 | IRAK3 | CSK |
| 19 | GRK2 | CAMK2B | | MAPK10 | | WNK3 | MAP3K12 | RIPK2 | FGFR1 |
| 20 | RPS6KA1_2 | DAPK3 | | MAPK13 | | AURKA | STK3 | KSR2 | EPHA8 |
| 21 | RPS6KA3 | CASK | | CDK9 | | WNK1 | MAP3K7 | KSR1 | LCK |
| 22 | PKN1 | PASK | | CDK1 | | WEE1 | MAP3K8 | LRRK2 | TYK2_2 |
| 23 | GRK5 | MARK2 | | MAPK7 | | PKMYT1 | MAP4K3 | | MST1R |
| 24 | GRK4 | MAPKAPK3 | | CDK16 | | WEE2 | MAP3K5 | | ERBB2 |
| 25 | PRKCQ | CAMK1 | | CDK8 | | PRAG1 | TAOK3 | | ITK |
| 26 | CDC42BPB | PIM2 | | DYRK3 | | CAMKK1 | PAK3 | | IGF1R |
| 27 | PRKG1 | PRKAA2 | | CLK2 | | EIF2AK2 | STK10 | | BMX |
| 28 | STK38 | MARK4 | | CLK4 | | ULK3 | MAP3K20 | | EPHB4 |
| 29 | RPS6KA6 | DCLK1 | | DYRK2 | | AURKC | TNIK | | EPHB2 |
| 30 | ROCK2 | STK40 | | PRPF4B | | TTK | STK4 | | MET |
| 31 | | MELK | | MAPK6 | | ULK1 | MAP2K7 | | ROR2 |
| 32 | | MKNK1 | | CLK1 | | ULK2 | MAP4K1 | | NTRK2 |
| 33 | | SNRK | | CDK7 | | ULK4 | | | SYK |
| 34 | | MARK1 | | HIPK2 | | ERN1 | | | JAK1 |
| 35 | | PRKAA1 | | CDK13 | | TP53RK | | | FGFR4 |
| 36 | | STK17B | | HIPK3 | | CSNK2A2 | | | ROS1 |

| | | | | | | | | | |
|---|---|---|---|---|---|---|---|---|---|
| 37 | | STK17A | | CDK4 | | PIK3R4 | | | EGFR |
| 38 | | TRIB2 | | CDK11B | | MLKL | | | NTRK3 |
| 39 | | | | | | CAMKK2 | | | JAK1_2 |
| 40 | | | | | | | | | EPHB3 |
| 41 | | | | | | | | | JAK3_2 |
| 42 | | | | | | | | | EPHB1 |
| 43 | | | | | | | | | BTK |
| 44 | | | | | | | | | JAK2_2 |
| 45 | | | | | | | | | AXL |
| 46 | | | | | | | | | JAK2 |
| 47 | | | | | | | | | PTK6 |
| 48 | | | | | | | | | NTRK1 |
| 49 | | | | | | | | | DDR2 |
| 50 | | | | | | | | | DDR1 |
| 51 | | | | | | | | | LMTK3 |
| 52 | | | | | | | | | ROR1 |
| 53 | | | | | | | | | RYK |
| 54 | | | | | | | | | PTK7 |
| 55 | | | | | | | | | ABL1 |
| 56 | | | | | | | | | MERTK |
| 57 | | | | | | | | | TYK2 |
| 58 | | | | | | | | | ALK |
| 59 | | | | | | | | | FGFR3 |
| 60 | | | | | | | | | RET |
| 61 | | | | | | | | | EPHA2 |

**Table S5** Allosteric residue index of kinases. In each of these representative structures, the kinase catalytic domain is renumbered from 1. The residue indices are shown in ascending order in the main text Fig. 5c-5i and supplementary Figures S8, S9.

| Site | Name of the Site | Representative Kinase and PDB id | Residue Number |
|---|---|---|---|
| Ortho | Orthosteric | ABL1 (PDB id: 1OPL) | 7, 8, 12, 15, 28-30, 45, 49, 58, 72-80, 129, 139-141 |
| MT3 | MEK1/2 type III inhibitor | MAP2K1 (PDB id: 4AN2) | 11, 30, 32, 48, 51, 60, 61, 74, 76, 123, 125, 128, 140-145, 148, 149, 152, 156-159 |
| AAS | Aurora A activation segment | AURKA (PDB id: 4C3P) | 20, 22, 23, 25, 27, 34, 37, 38, 43, 46, 47, 50, 51, 54, 55, 65-69, 74, 77 |
| PDIG | PDIG motif | CHEK1 (PDB id: 3JVS) | 85-90, 125, 126, 165, 192, 196-198 |
| PIF | PDK1 interacting fragment | PDPK1 (PDB id: 4RQK) | 34, 37, 38, 43, 46, 47, 50, 67-69, 74-76 |
| CMP | c-Abl myristoyl pocket | ABL1 (PDB id: 1OPL) | 96, 99, 100, 103, 188, 191, 192, 194, 221-224, 227 |
| MPP | MKK4 p38a peptide | MAP2K4 (PDB id: 3ALO) | 1, 20, 25, 27, 41, 45, 62-69, 76 |
| DRS | D-recruitment site | MAPK8 (PDB id: 1UKI) | 87, 93, 96, 101, 102, 105, 106, 108, 134-138 |
